# Homeodomain protein Sxi1α independently controls cell-cell fusion and gene expression during sexual reproduction in *Cryptococcus deneoformans*

**DOI:** 10.1101/2025.02.11.637763

**Authors:** Jun Huang, Anna E. Lehmann, Patricia P. Peterson, Ziyan Xu, Liping Xiong, Sheng Sun, Joseph Heitman

**Author notes:** Corresponding author: Joseph Heitman.

## Abstract

Sex-specific homeodomain (HD) proteins are key regulators of cell identity and sexual development in fungi, typically functioning as heterodimers to govern transcription. In the human fungal pathogens *Cryptococcus neoformans* and *Cryptococcus deneoformans*, the HD proteins Sxi1α and Sxi2**a** (Sex-inducer 1α and 2**a**) have been characterized as interacting components that play critical roles in sexual development during α x **a** sexual reproduction. α cells are the predominant mating type in natural populations of *Cryptococcus*, and unisexual (same-sex) mating can also occur in certain genetic backgrounds. The roles of Sxi1α and Sxi2**a** in unisexual reproduction are not fully understood. To elucidate the functions of Sxi1α and Sxi2**a**, we first applied AlphaFold3 prediction, which identified potential heterodimeric and homodimeric complexes. Formation of a Sxi2**a** homodimer was then experimentally validated through yeast two-hybrid assays. We subsequently deleted *SXI1*α and *SXI2***a** in the hyper-filamentous self-fertile *C. deneoformans* strains XL280α and XL280**a**. Disruption of these genes did not result in noticeable defects in vegetative growth, virulence-associated traits, colony morphology, sporulation, or competitive fitness during unisexual crosses. Interestingly, both bilateral (mutant x mutant) and unilateral (mutant x wildtype) crosses involving the *sxi1*αΔ mutant significantly increased α-α cell fusion efficiency, suggesting a previously unrecognized inhibitory role for Sxi1α in regulating same-sex cell fusion. Consistently, genes encoding mating pheromones and the α-pheromone receptor Ste3 were upregulated in the *sxi1*αΔ fusion assays. Transcriptomic analysis of *sxi1*αΔ and *sxi2***a**Δ mutants led to the identification of unique subsets of genes negatively regulated by each transcription factor during unisexual reproduction. Additionally, α x **a** crosses between null mutants of *sxi1*αΔ and *sxi2***a**Δ revealed differential regulation of mating-type (*MAT*) loci genes dependent only on Sxi1α or Sxi2**a**. Together, our findings reveal a novel role for Sxi1α in governing cell fusion and demonstrate that Sxi1α and Sxi2**a** have distinct transcriptional control during unisexual and α x **a** sexual reproduction, potentially exerting opposing regulation of sex-specific *MAT* genes.

**Author Summary:** Analogous to processes observed in many eukaryotic organisms, fungal sexual reproduction reshuffles genetic material, producing novel genetic variants with increased fitness and/or adaptive potential. The human fungal pathogen *Cryptococcus deneoformans* is widespread in the environment and can cause severe infections in immunocompromised individuals. Sexual reproduction in this species occurs between cells of either opposite mating types (α x **a** mating) or the same mating type (α x α and **a** x **a** unisexual reproduction). The mating type-specific homeodomain transcription factors Sxi1α and Sxi2**a** are known to form a complex that broadly controls sexual development following α and **a** cell fusion. Here, we find that Sxi1α inhibits cell-cell fusion during unisexual reproduction, while its partner, Sxi2**a**, shows only limited inhibition in certain genetic contexts. Further, we show that Sxi1α and Sxi2**a** regulate transcription of distinct gene sets during both α x **a** and unisexual reproduction, including many within the mating-type (*MAT*) loci, often in opposing manners. Together, these findings reveal that Sxi1α and Sxi2**a** function both cooperatively and independently, thereby allowing for a more flexible and finely-tuned mating control system than previously recognized, providing insight into how this organism generates genetic variation and adapts to changing environments.

## Introduction

Sexual reproduction is a key process in eukaryotic organisms, promoting genetic diversity and facilitating adaptation to changing environments. In fungi, this process is controlled by the mating-type (*MAT*) locus, a specific genomic region containing genes required for successful mating [1]. Among these genes, pheromone and pheromone receptors (P/R) are involved in signal transduction and recognition between mating partners, and homeodomain (HD) transcription factors act as master regulators that control the expression of genes necessary for sexual development [2,3]. In the model yeast *Saccharomyces cerevisiae*, **a**1 and α2 are well-characterized HD proteins, forming **a**1-α2 heterodimers that bind DNA cooperatively to repress transcription of cell type-specific genes [4,5]. Unlike **a**1, which has no known role in the absence of α2, α2 can form homodimers, as well as partner with the Mcm1 cofactor to repress **a**-specific genes in α and **a**/α cells [6–9]. In the corn smut fungus *Ustilago maydis*, the HD proteins bWest (bW) and bEast (bE) dimerize to regulate fungal morphological switching and interactions with the plant host [10–13]. Similarly, sexual development in the mushroom *Coprinus cinereus* requires the physical interaction of two HD proteins, HD1 and HD2 [14]. Conserved and convergent functionality of HD transcription factors across diverse fungi illustrates their centrality to species survival and proliferation.

The human fungal pathogen *Cryptococcus neoformans* is the predominant cause of cryptococcal meningoencephalitis worldwide, whereas its sister species *C. deneoformans* displays a more geographically restricted distribution and is more frequently encountered in certain regions, particularly in Europe [15–17]. Interestingly, in *C. neoformans* and *C. deneoformans*, the *P/R* and *HD* genes as well as additional genes reside in a relatively large (>100 kb) *MAT* locus [2,17–19]. In addition to its direct role in α x **a** sexual reproduction, the *MAT* locus is associated with several other functions in *Cryptococcus*, including virulence and uniparental mitochondrial inheritance (mito-UPI), in which mitochondrial DNA is predominantly inherited from the *MAT***a** parent after α x **a** sexual reproduction [20,21]. We previously identified the *Cryptococcus* sex-specific HD proteins Sxi1α (Sex inducer 1α) and Sxi2**a** (Sex inducer 2**a**), which heterodimerize upon α-**a** cell fusion and are involved in key development processes including cell identity determination, sexual development, and mito-UPI [2,21,21–25]. Our group and others have identified several Sxi1α-Sxi2**a** targets for which proper temporal expression is essential for sexual development. For example, *CPR2* (*Cryptococcus* pheromone receptor 2) governs α**-a** cell fusion, and *CLP1* (clampless 1) is required for post-cell fusion dikaryotic filament formation [26–29]. Beyond these well-characterized targets, the identification of several hundred Sxi1α/Sxi2**a**-bound promoter elements in *C. neoformans* suggests that this complex has a much broader regulatory role in sexual development [29].

In addition to α x **a** sexual reproduction, unisexual reproduction can also occur between partners of the same mating type (mainly α x α) in *C. neoformans* and *C. deneoformans* [30–32]. In natural *Cryptococcus* populations, *MAT*α isolates are predominant among both clinical and environmental samples, suggesting that α x **a** heterothallic sexual reproduction is infrequent [20,33], even though population genomics studies have shown that recombination is occurring among *Cryptococcus* isolates in nature [33,34]. This strong bias in *MAT* locus distribution may, at least in part, be explained by the occurrence of unisexual reproduction, which enables genetic recombination and propagation in the absence of a compatible mating partner [30,35,36]. As key sex-specific transcriptional regulators, the roles of Sxi1α and Sxi2**a** in unisexual reproduction and their transcriptional targets remain to be elucidated.

In this study, we sought to determine whether Sxi1α and Sxi2**a** are involved in same-sex mating in *C*. *deneoformans*. AlphaFold3 structural prediction suggested that Sxi1α and Sxi2**a** may form both heterodimeric and homodimeric complexes, and the formation of the Sxi2**a** homodimer was further confirmed by yeast two-hybrid assays. Taking advantage of the hyper-filamentous *C. deneoformans* congenic isolates XL280α and XL280**a** [37], along with a CRISPR/Cas9-based gene editing system [38,39], we deleted *SXI1*α and *SXI2***a** in XL280α and XL280**a**, respectively. Consistent with previous results, we found that *SXI1*α and *SXI2***a** are dispensable for vegetative growth and multiple *in vitro* virulence-relevant traits, including thermotolerance at 37°C, polysaccharide capsule formation, and melanin production. Interestingly, compared to fusion between two wildtype *MAT*α strains, we observed a significant increase in α-α fusion efficiency when Sxi1α was deleted in one or both *MAT*α strains, which is accompanied by significant upregulation of genes encoding mating pheromones and the α-pheromone receptor *STE3*. Similarly, increased **a**-**a** fusion was observed when Sxi2**a** was deleted in one of the two *MAT***a** strains, but not when Sxi2 was absent in both strains, pointing to a previously unrecognized inhibitory function of Sxi1α in same-sex cell fusion. No apparent defects in colony morphology, hyphal development, basidium formation, sporulation, spore germination, or competitive fitness were detected in *sxi1*αΔ or *sxi2***a**Δ mutants during unisexual reproduction. Transcriptomic analysis revealed distinct sets of genes for which expression differences are regulated only by Sxi1α, independent of its canonical interacting partner Sxi2**a**. The Sxi1α-dependent gene set in the unisexual crosses is markedly smaller and has limited overlap with that in α x **a** crosses. These observations support the hypothesis that Sxi1α has distinct functions during α x **a** and α x α sexual reproduction that is independent of Sxi2**a**.

Our results suggest a novel function of Sxi1α in regulating cell-cell fusion, expanding our understanding of how HD proteins contribute to fungal sexual reproduction. Additionally, we show that Sxi1α and Sxi2**a** have transcriptional regulatory effects both dependently and independently of one another. These findings, combined with our structural predictions and yeast two-hybrid-validated Sxi2**a** homodimer, support the hypothesis that Sxi1α and Sxi2**a** may have distinct functions independently of their usual HD partner, analogous to α2 in budding yeast. Overall, these findings further our understanding of how sexual reproduction is regulated in this important human fungal pathogen and reveal important differences in Sxi1α and Sxi2**a** transcription factor activity between unisexual and α x **a** sexual reproduction.

## Results

### Sxi1α and Sxi2a are predicted to form a heterodimer

To study the impact of the homeodomain proteins Sxi1α and Sxi2**a** on unisexual reproduction in *Cryptococcus*, we selected to study XL280α and XL280**a**, a pair of congenic *C. deneoformans* strains that are well-known for their ability to undergo unisexual reproduction involving robust self-filamentation and sporulation [30,37]. Sxi1α and Sxi2**a** are conserved between *C. neoformans* and *C. deneoformans*, and homeodomain (Pfam:00046) or homeobox KN domains (from **KN**OX family transcription factors, Pfam:PF05920) were found within the sequences via an NCBI conserved domain search (Fig 1A). Both domains are structurally related, sharing DNA-binding functions through a helix-turn-helix motif, consistent with potential functional relevance in governing gene expression. Consistent with our previous report with yeast two-hybrid analysis [2,22], a heterodimer between Sxi1α and Sxi2**a** was predicted by AlphaFold3 protein-protein complex prediction (Fig 1B). Pairwise structural alignments between the predicted Sxi1α-Sxi2**a** heterodimer and the crystal structure of the *S. cerevisiae* **a**1-α2 heterodimer [40] revealed robust structural similarity between the two homeodomain complexes with an RMSD value of 0.561 angstroms between 49 pruned atom pairs (Fig 1C). Similarly, the Sxi1α-Sxi2**a** heterodimer exhibited high structural similarity to the AlphaFold3-predicted bW-bE heterodimer from the corn smut fungus *Ustilago maydis*, with a RMSD of 1.069 angstroms across 40 pruned atom pairs. Interestingly, the HD2 proteins [2], represented by Sxi2**a**, **a**1, and bW, superimpose closely, reflecting strong evolutionary conservation. In contrast, greater structural divergence was observed among the HD1 proteins, including Sxi1α, α2, and bE, consistent with their roles in conferring regulatory specificity. Together, these comparisons support a conserved functional framework in which HD2 provides a structurally stable DNA-binding platform, while HD1 contributes specificity to transcriptional regulation, with heterodimer formation serving as a central regulatory module in mating control [41].

**Figure 1.**
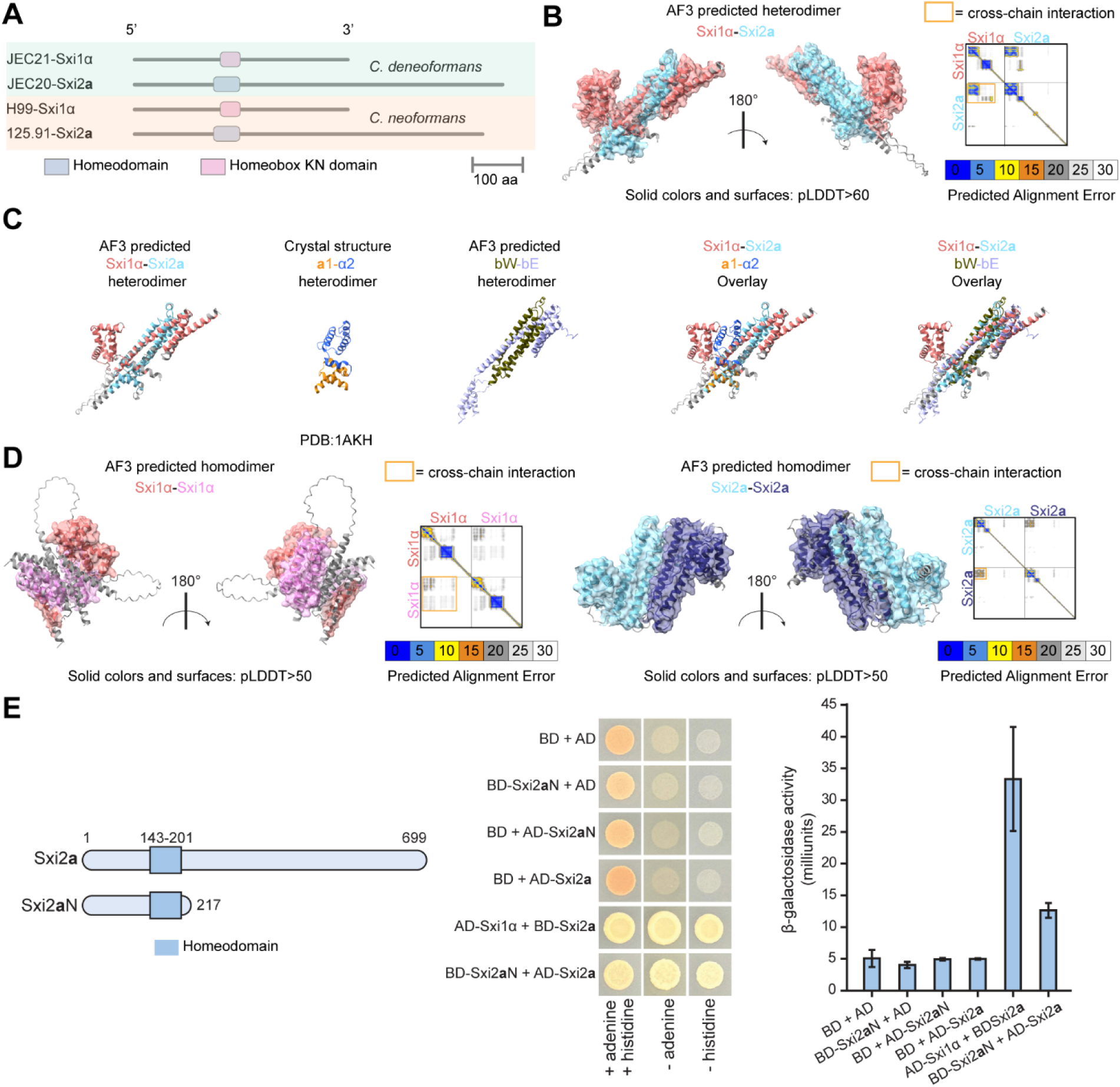
AlphaFold3 predicts heterodimer and homodimers of Sxi1α and Sxi2a. (A) Predicted domain structures of Sxi1α and Sxi2**a** in *C. neoformans* and *C. deneoformans*. (B) AlphaFold3 (AF3)-predicted heterodimer structure of Sxi1α and Sxi2**a** from *C. deneoformans*. C-terminal disordered regions were hidden for visualization. Only regions with a predicted local distance difference test (pLDDT) score >60 were colored and surfaced. The predicted alignment error (PAE) plot of the predicted Sxi1α-Sxi2**a** heterodimer is displayed in the right panel. (C) Overlay of AlphaFold3-predicted Sxi1α-Sxi2**a** heterodimer with crystal structure of **a**1-α2 (from *S. cerevisiae*, PDB:1AKH) and predicted bW-bE heterodimer (from *U. maydis*, with high confidence ipTM = 0.74 pTM = 0.64). Pairing was performed with the “Matchmaker” function in Chimera X. (D) AlphaFold3-predicted homodimer structures with PAE plots for Sxi1α-Sxi1α (left panel) and Sxi2**a**-Sxi2**a** (right panel). Only regions with a pLDDT score >50 were colored and surfaced. (E) Yeast two-hybrid analysis of interactions among Sxi1α, Sxi2**a**, and the N terminal region of Sxi2**a** (Sxi2**a**N). Schematics of Sxi2**a** and Sxi2**a**N constructs are shown on the left, with the homeodomain indicated. *S. cerevisiae* strains expressing Gal4 DNA-binding domain (BD) or Gal4 activation domain (AD) fusions were mated, and diploids were selected on synthetic dextrose medium. Protein-protein interactions between bait and prey fusions drive Gal4-dependent activation of *ADE2* and *HIS3* reporter genes, allowing for growth on medium lacking adenine or histidine. Activation of *ADE2* in the *ade2* mutant reporter strain background also reduces red pigment accumulation, resulting in lighter colony coloration on nonselective medium. Quantification of *lacZ* reporter activation was performed using a chlorophenol red-β-D-galactopyranoside (CPRG) liquid assay, and β-galactosidase activity is shown on the right. Error bars indicate mean ± standard deviation from three technical replicates.

Previous studies showed that, in addition to forming a heterodimer with **a**1, *S. cerevisiae* α2 forms a homodimer to repress **a**-specific genes [9]. To benchmark the reliability of our structural predictions, we applied AlphaFold3 to the well-characterized yeast **a**1 and α2. Consistent with experimental evidence, AlphaFold3 predicted robust homodimer formation for α2 but not for **a**1, supporting the validity of this approach for distinguishing biologically relevant homodimerization events (S1 Fig). Using this validated framework, we next asked whether Sxi1α and Sxi2**a** might also form homodimers, we utilized AlphaFold3 to model Sxi1α-Sxi1α and Sxi2**a**-Sxi2**a** interactions. The predictions indicated potential low-confidence homodimer formation for Sxi1α-Sxi1α (ipTM = 0.14, pTM = 0.23) and Sxi2**a**-Sxi2**a** (ipTM = 0.20, pTM = 0.24) (Fig 1D). These low interface and predicted TM-scores suggest weak or unstable interactions. It is also possible that any homodimerization could involve additional interacting partner(s) that are not yet known, or that Sxi1α and Sxi2**a** could have independent roles analogous in function but distinct in structure from those observed in ascomycetous yeast.

To determine whether Sxi1α or Sxi2**a** could form homodimers, we performed yeast two-hybrid (Y2H) analyses with Gal4 DNA-binding domain (GBD) and transcriptional activation domain (GAD) fusion constructs (Fig 1E). Because homeodomain transcription factors frequently function as dimers, we specifically tested whether either protein could self-associate in this heterologous system. We also included the putative *C. deneoformans* ortholog of Mcm1, a transcription factor involved in the pheromone response pathway in other fungi including *S. cerevisiae*, where it interacts with the homeodomain protein α2 and transcription factor Ste12 to regulate mating-specific gene expression [42,43]. We identified C*. deneoformans* Mcm1 (CNN00060) and *C. neoformans* Mcm1 (CNAG_07924) as orthologs of *S. cerevisiae* Mcm1 through reciprocal BLASTp searches. Our AlphaFold3 prediction and alignment suggest a conserved structural organization between the *S. cerevisiae* α2-Mcm1 [44] and the *C. deneoformans* Sxi1α-Mcm1(CNN00060) (S2 Fig), and we hypothesize that Mcm1 could similarly interact with Sxi1α or Sxi2**a** in *Cryptococcus*. Thus, we tested all possible pairwise interactions between Sxi1α, Sxi2**a**, and Mcm1.

Plasmids encoding the fusion proteins were transformed into the Y2H reporter strains Y2HGold and Y187. Following mating of the two strains, diploids were selected and assayed for Gal4-dependent activation of *ADE2*, *HIS3*, and *lacZ* reporter genes. In a previous study, full-length Sxi2**a** fused to GBD was shown to auto-activate reporter gene expression [2]. Consistent with this observation, auto-activation was also detected in our assays for both Sxi2**a** and Mcm1 (S3 Fig). To avoid false-positive interactions resulting from auto-activation, we instead analyzed a truncated form of Sxi2**a** (Sxi2**a**N), containing only the N-terminal region and the homeodomain (HD), for interaction testing. A robust protein-protein interaction was detected between Sxi2**a**N and full-length Sxi2**a**, supporting the AlphaFold-predicted model of a Sxi2**a** homodimer (Fig 1E). In contrast, we did not detect evidence for Sxi1α homodimerization, or between Mcm1 and Sxi1α or Sxi2**a** in this assay (S4 Fig). While these results suggest that such interactions are either weak or absent in this context, we cannot exclude the possibility that dimerization or complex formation may occur *in vivo* and require post-translational modifications or additional binding partners not recapitulated in the yeast two-hybrid system.

### Sxi1α and Sxi2a are dispensable for *in vitro* growth and virulence-related traits

To further characterize the phenotypic impact of Sxi1α and Sxi2**a**, we generated *SXI1*α and *SXI2***a** null mutants in XL280α and XL280**a**, respectively, with a CRISPR/Cas9-based strategy following the previously-established TRACE electroporation protocol [38,39]. Multiple independent drug-resistant transformants [four for XL280α *sxi1*αΔ::*NAT* and two for XL280**a** *sxi2***a**Δ::*NEO*] were constructed and confirmed with correct genotypes as deletion mutants based on 5’ junction, 3’ junction, and internal ORF-specific PCR analysis (Fig 2A). Mapping of reads generated from Illumina whole-genome sequencing further confirmed the complete deletion of the genes in all four *sxi1*αΔ mutants and one of the two *sxi2***a**Δ mutants, and a partial gene deletion in the other *sxi2***a**Δ mutant (Fig 2B and S5 Fig). In some of the mutants, elevated read-depth was observed in the flanking regions employed for homologous recombination during gene deletion (Fig 2B and S5 Fig). This could indicate potential concatemeric or ectopic insertion [45] of the resistance cassette (Fig 2B and S5 Fig). For the partial deletion mutant XL280**a** *sxi2***a**Δ#6, the deletion occurred between the 5’ microhomology sequence and the second gRNA targeting site, instead of at the 3’ microhomology sequence (S5 Fig). This pattern is consistent with a hybrid integration event in which NHEJ and HR [46] occurred on opposite sides of the construct. Importantly, no sequencing reads mapped to the N-terminal region of the gene that encompasses the conserved homeodomain (S5 Fig). Therefore, Sxi2**a** is functionally disrupted in this mutant, and the strain was retained for subsequent analyses.

**Figure 2.**
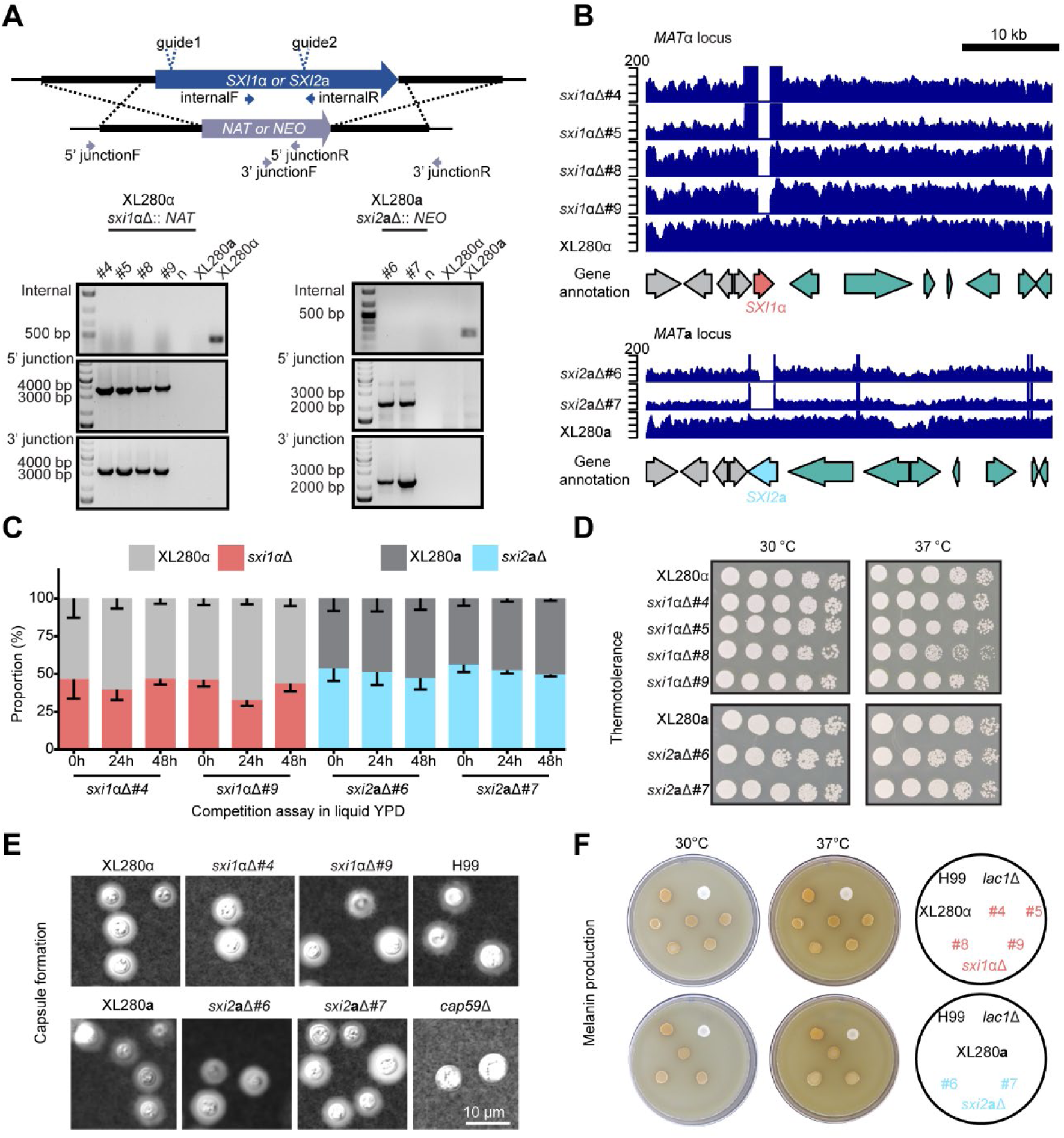
Sxi1α and Sxi2a are not involved in growth, capsule formation, and melanin production. (A) Generation of independent *sxi1*αΔ and *sxi2***a**Δ mutants in XL280α and XL280**a**, respectively. Transformants were verified through internal PCR targeting the ORF, along with 5’ and 3’ junction PCRs specific to the drug resistance marker to confirm proper integration at the genomic locus. (B) Whole-genome sequencing (WGS) reads mapping confirming the complete or partial deletion of *SXI1*α (red block arrow) and *SXI2***a** (blue block arrow). Genes located within and outside of the mating-type locus are highlighted in green and gray, respectively. (C) *In vitro* competition assays of the *sxi1*αΔ and *sxi2***a**Δ mutants, each carrying either the *NAT* or *NEO* resistance markers, and their corresponding XL280 wildtype strains. Each competition was assayed after 0, 24, 48 hours of incubation, with four independent replicates. Error bars indicate standard deviation. (D) Serial dilutions of *sxi1*αΔ and *sxi2***a**Δ mutants, along with the corresponding XL280 wildtype strains, were spotted on solid YPD media and incubated at 30°C and 37°C. (E) Saturated cultures of the indicated strains, grown in RPMI medium, were stained with India ink to visualize capsule formation. The H99 strain and isogenic *cap59*Δ mutant served as positive and negative controls, respectively. (F) Overnight cultures of the indicated strains were spotted on Niger seed medium to monitor melanin formation. The wildtype H99 strain and isogenic *lac1*Δ mutant served as positive and negative controls, respectively.

We next performed competition assays to assess the impact of Sxi1α and Sxi2**a** on fungal fitness during vegetative growth. Equal amounts of mutant and wildtype cells were inoculated in liquid YPD medium and cultured for 48 hours at 30°C. Samples were collected at 0h, 24h, and 48h to assess population dynamics. In all cases, the *sxi1*αΔ and *sxi2***a**Δ strains maintained similar proportions to wildtype during co-culture, indicating that the mutations confer no detectable fitness defect (Fig 2C). Additionally, we characterized the contributions of Sxi1α and Sxi2**a** to several major virulence traits: thermotolerance at 37°C, capsule formation, and melanin production. Consistent with our previous reports in *C. neoformans* [23], none of the *sxi1*αΔ or *sxi2***a**Δ *C. deneoformans* deletion strains displayed fitness differences compared to the XL280 wildtype strains in these assays (Fig 2D-2F). Taken together, these results indicate that the loss of *SXI1*α and *SXI2***a** does not affect the fitness of the XL280 strain during vegetative growth and has no detectable impact on virulence-related traits under the conditions tested.

### Sxi1α and Sxi2a are dispensable for morphology, sporulation, and cell viability during unisexual reproduction

We next investigated whether Sxi1α and Sxi2**a** are involved in unisexual reproduction in the XL280 background. When grown on mating- and selfing-inducing MS solid medium [47] as solo-cultures, all of the tested *sxi1*αΔ and *sxi2***a**Δ mutants produce elongated hyphae, basidia, and basidiospore chains at levels similar to the XL280 wildtype controls (Fig 3A and 3B). No discernable differences in the basidia and basidiospores were observed by scanning electron microscope (SEM) (Fig 3C). Thus, the deletion of *SXI1*α and *SXI2***a** did not affect unisexual reproduction morphologically in the XL280 background. Additionally, we collected random meiotic basidiospores by microdissection to test the effect of Sxi1α and Sxi2**a** on spore germination rates and cell viability and found no significant difference in germination rates between spores produced by the *sxi1*αΔ mutants and XL280α (p-values: 0.682 (*sxi1*αΔ#4 vs XL280α) and 0.125 (*sxi1*αΔ#5 vs XL280α), Welch’s t-test), as well as between spores produced by the *sxi2***a**Δ mutants and XL280**a** (p-values: 0.591 (*sxi2***a**Δ#6 vs XL280**a**:) and 0.750 (*sxi2***a**Δ #7 vs XL280**a**), Welch’s t-test) (Fig 3D). No significant difference in spore germination rate was observed between the congenic wildtype strains XL280α and XL280a (p-value: 0.0856, Welch’s t-test) (Fig 3D).

**Figure 3.**
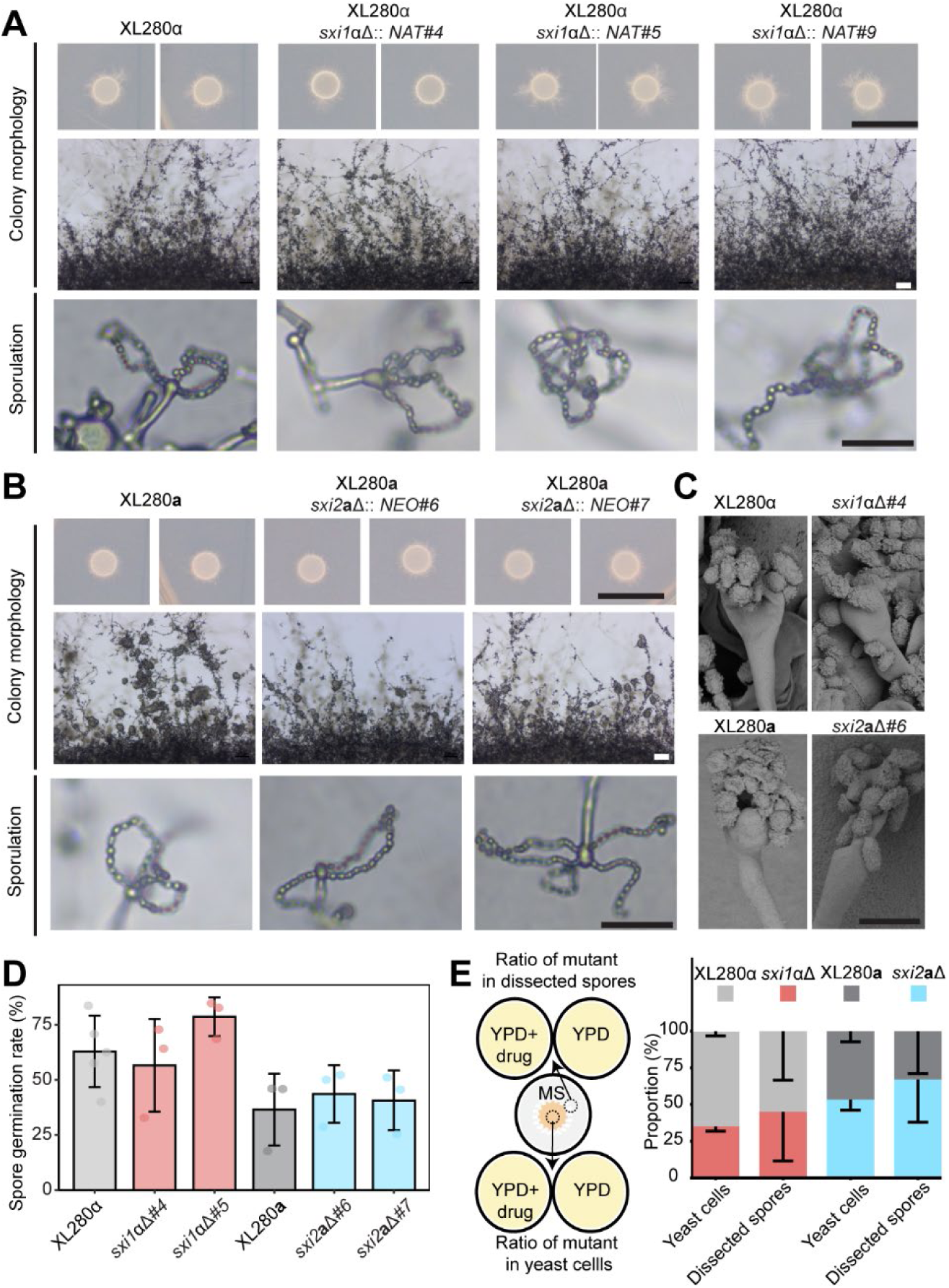
Sxi1α and Sxi2a are not required for hyphal growth or basidium and spore production. (A and B) Colony morphology (top), self-filamentation (middle), and sporulation (bottom) phenotypes of independent *sxi1*αΔ (A) and *sxi2***a**Δ (B) mutants when grown under mating-inducing conditions. Scale bars, 1 cm (upper panel), 100 μm (middle panel) and 20 μm (bottom panel). (C) Scanning electron microscopy (SEM) images of basidia and basidiospores from the indicated solo-culture. Samples were grown on MS media for 1 to 2 weeks before microscopy. Scale bar: 5 μm. (D) Germination rate of random spores dissected from the indicated solo-cultures. At least 3 replicates were performed for each strain (indicated by dots), with error bars depicting standard deviation among replicates. (E) Competition assays of *sxi1*αΔ, *sxi2***a**Δ mutants against their corresponding XL280 wildtype strain. The co-cultures were grown on MS medium under mating-inducing conditions. Spores from the periphery, and yeast cells from the center, of the co-culture spots were dissected and analyzed to determine their origin (i.e., wildtype or mutant), which were then used to infer the relative competitiveness of the mutant strains. Five different co-culture spots were analyzed for each competition.

We previously observed that strains capable of undergoing unisexual reproduction outcompete mutants with unisexual reproduction defects under mating-inducing conditions, attributable to enhanced hyphal growth and sporulation [48]. To assess the effects of *SXI1*α and *SXI2***a** deletion on this metric of unisexual reproduction efficiency, for each deletion strain we mixed equal numbers of cells from the mutant (containing a dominant drug-resistant marker) and its wildtype control strain (sensitive to drug), and spotted the mixture on MS medium to induce unisexual reproduction. After robust sporulation was observed (10 days post plating), we dissected two populations: 1) random basidiospores from the periphery where sporulation occurs, and 2) the center of the spot where cells remain mostly as yeast (Fig 3E, left panel). Germinated dissected spores/yeast cells were transferred onto YPD plates containing selective drug (NAT or NEO) to determine which strain in the mixture they originated from. For both *sxi1*αΔ and *sxi2***a**Δ, the ratios between mutant and wildtype among the basidiospores at the periphery were similar to those among the yeast cells at the center (spores/yeast, p-values: 0.5398275 (*sxi1*αΔ) and 0.379667 (*sxi2***a**Δ), paired t-test), indicating the loss of Sxi1α and Sxi2**a** does not affect competitive fitness during unisexual reproduction (Fig 3E, right panel).

The DNA-binding protein Mcm1 functions as a central transcriptional co-regulator of mating type homeodomain proteins during sexual development in budding yeast [43,44]. Because *Cryptococcus* Sxi1α and Sxi2**a** have structural and functional commonalities with *S. cerevisiae* α2 and **a**1, we questioned whether an analogous co-factor may exist in *Cryptococcus*. We first examined whether deletion of *MCM1* affects unisexual reproduction by generating *mcm1*Δ mutants in the XL280 background (S6 Fig) and tested their self-fertility under mating-inducing conditions. Deletion of *MCM1* did not result in obvious defects in unisexual mating morphology in either XL280α or XL280**a** (S7 Fig). To test for potential synthetic effects between *MCM1* and *SXI1*α or *SXI*2**a**, we constructed *sxi1*αΔ *mcm1*Δ and *sxi2*aΔ *mcm1*Δ double mutants (S6 Fig). No clear unisexual mating morphology defects were observed in these strains (S7 Fig). Interestingly, it appears that all of the engineered strains containing *mcm1*Δ deletion displayed an increased abundance of yeast cells in the center of mating spots, along with a reduced frequency of basidia bearing robust basidiospore chains (S7 Fig). Additionally, we tested the role of Mcm1 in α x **a** sexual reproduction with an *mcm1*Δ strain derived from the *C. neoformans* H99 transcription factor deletion collection [49]. Through mating, we generated the congenic strain KN99**a** *mcm1*Δ. Crosses were performed on MS and V8 (pH=5) media by mating *mcm1*Δ with either wildtype, *sxi1α*Δ, or *sxi2**a***Δ strains [23]. Consistent with previous observations [2,22], crosses involving *sxi1α*Δ or *sxi2a*Δ exhibited clear mating defects in hyphal development and sporulation (S8 Fig). In contrast, no detectable mating defects were observed in either unilateral or bilateral α x **a** crosses involving *mcm1*Δ (S8 Fig). Collectively, these results indicate that, unlike its essential co-regulatory role in budding yeast, Mcm1 is largely dispensable for both unisexual and α x **a** sexual reproduction in *C. deneoformans* and *C. neoformans*.

### Sxi1α is critical for unisexual cell-cell fusion

In the presence of a helper strain serving as a pheromone donor, α x α and **a** x **a** outcrossing can be facilitated in *C. deneoformans* in so-called Ménage à trois matings [22,30] (Fig 4A). Previously, our results from α x **a** sexual reproduction suggested that Sxi1α and Sxi2**a** function after cell-cell fusion [2]. Although cell-cell fusion is dispensable for solo unisexual reproduction, it may still occur under certain conditions, such as during α x α outcrossing [22,30]. Given the limited understanding of the role of Sxi1α in this context, we sought to determine whether Sxi1α is involved in cell-cell fusion during α x α outcrossing in the presence of a donor strain (JEC20, *MAT***a** *NAT^S^ NEO^S^*). Strains in the XL280α *SXI1*α and *sxi1*αΔ backgrounds that were differentially resistant to *NAT* and *NEO* were used to set up three types of mating: *SXI1*α x *SXI1*α, *SXI1*α x *sxi1*αΔ, and *sxi1*αΔ x *sxi1*αΔ. Equal number of cells of the two strains in each mating were mixed and spotted on V8 agar (pH=7) to induce sexual reproduction. After 72 hours of incubation, cells were collected. The fusion frequency was fewer than 5.04x10^-8^ in two independent sets of the XL280α *NAT* x XL280α *NEO* crosses, with fusion products identified by screening colonies for NAT and NEO resistance. In contrast, significantly higher fusion frequencies were found in *sxi1*αΔ::*NAT* x *sxi1*αΔ::*NEO* (22.5-fold higher than wildtype, frequency: 1.13x10^-6^, p-value: 0.0013, Welch’s t-test) and *sxi1*αΔ::*NAT* x XL280α *NEO* (62.9-fold higher than wildtype, frequency: 3.17x10^-6^, p-value: 0.0288, Welch’s t-test) (Fig 4B). Taken together, these findings demonstrate that cell-cell fusion between *MAT*α cells is significantly increased when *SXI1*α is absent in at least one of the two strains.

**Figure 4.**
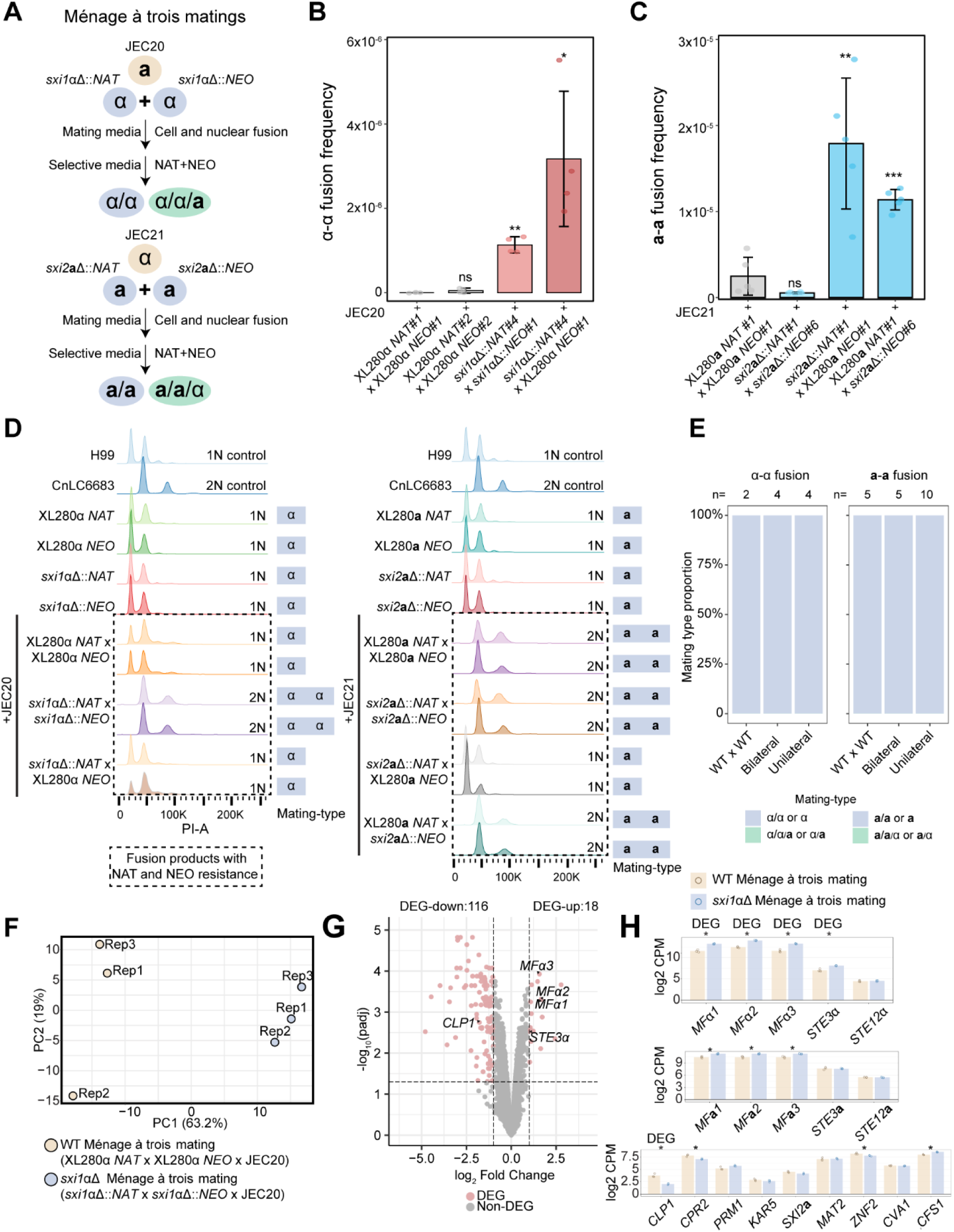
Fusion frequency and ploidy analysis of products from α x α and a x a Ménage à trois matings. (A) Schematic of the Ménage à trois mating assay. In each mating-type background, two strains carrying different selectable markers (NAT or NEO) were co-cultured with a pheromone donor of the opposite mating type to induce cell-cell fusion. NAT and NEO resistant colonies were recovered on selective medium. (B-C) Quantification of α x α (B) and **a** x **a** (C) fusion frequencies for the indicated bilateral and unilateral crosses. Cells from the indicated mating assay on V8 agar (pH = 7.0) for 72 h were plated on YPD + NAT + NEO and YPD to determine the ratio of cell-cell fusion products in the population. Data are presented as mean ± SD from four to five independent biological replicates; each point represents an individual replicate. ns, not significant; * (p-value < 0.05), ** (p-value < 0.01), *** (p-value < 0.001) indicate statistical significance compared to wildtype control. (D) Fluorescence-activated cell sorting (FACS) analysis of DNA content in parental strains and fusion products. H99 (1N control) and CnLC6683 (2N control) originate from the same experiment; their profiles are duplicated in both the left and right panels to facilitate comparison. Fusion products (highlighted by dashed boxes) are annotated with their mating type(s) as determined by genotyping PCR (right). 1N and 2N indicate haploid and diploid DNA content, respectively. (E) Proportion of mating type of fusion products from the indicated crosses. (F) Principal component analysis (PCA) of 3’ enriched RNA sequencing data from WT (XL280α *NAT* x XL280α *NEO* x JEC20) and *sxi1*αΔ bilateral (*sxi1*αΔ::*NAT* x *sxi1*αΔ::*NEO* x JEC20) Ménage à trois mating harvested after 72 h on V8 (pH=7) medium. Each point represents an independent biological replicate. The percentage of variance explained by PC1 and PC2 is indicated. (G) Volcano plot showing differential gene expression between *sxi1*αΔ bilateral and WT ménage à trois mating. Genes classified as differentially expressed are highlighted in red based on | log_2_ fold change| >=1 and adjusted p-value <= 0.05 thresholds, while non-differentially expressed genes are shown in gray. Selected genes of interest are labeled. (H) Bar plots showing normalized CPM expression levels for representative differentially expressed genes DEG and non-differentially expressed genes across WT and *sxi1*αΔ bilateral ménage à trois mating. Data are presented as mean ± standard deviation across biological replicates. Asterisks indicate statistically significant differences. Statistical significance was assessed using two sample Welch t tests on TMM normalized log_2_ CPM values for each gene, with p values adjusted for multiple testing with the Benjamini–Hochberg method. Genes labeled as DEG correspond to those classified as differentially expressed in the DEG analysis.

We then asked whether Sxi2**a** has a similar effect on cell-cell fusion during **a** x **a** unisexual mating (Fig 4A). Interestingly, with the presence of a donor strain (JEC21, *MAT*α, *NAT^S^ NEO^S^*), the fusion frequency between XL280**a** *NAT* and XL280**a** *NEO* was 2.43x10^-6^, while no significant difference in fusion frequency was found in *sxi2***a**Δ::*NAT* x *sxi2***a**Δ::*NEO* (20% of the wildtype, frequency 5.43x10^-7^, p-value: 0.1204, Welch’s t-test). In contrast, the fusion frequencies between *sxi2***a**Δ::*NAT* x XL280**a** *NEO* (7.2-fold higher than wildtype, frequency 1.79x10^-5^, p-value: 0.0086, Welch’s t-test), XL280**a** *NAT* x *sxi2***a**Δ::*NEO* (4.6-fold higher than wildtype, frequency 1.13x10^-5^, p-value: 0.0002, Welch’s t-test) were significantly higher than the wildtype cross (Fig 4C).

We hypothesized that NAT and NEO resistant colonies, carrying both selectable markers after cell-cell fusion event, would exhibit diploid DNA content, or triploid if the fusion involved a non-selective donor strain (Fig 4A). To verify the ploidy of fusion products from α x α and **a** x **a** outcrosses, flow cytometry analysis was performed. With *C. neoformans* haploid strain H99 and diploid strain CnLC6683 as controls [50], all NAT- or NEO- containing outcrossing parent strains were confirmed to possess haploid DNA content (Fig 4D). Two independent NAT and NEO resistant colonies from Ménage à trois matings were randomly selected for analysis. As expected in the α x α fusion assay, diploid DNA content was observed in bilateral crosses between *sxi1*αΔ deletion mutants. In contrast, haploid DNA content was detected in XL280α *NAT* x XL280α *NEO* and *sxi1*αΔ::*NAT* x XL280α *NEO* crosses. The presence of haploid NAT and NEO resistant colonies suggests that our assay may slightly overestimate the true frequency of cell-cell fusion events, likely due to genome recombination during unisexual reproduction. Similarly, in **a** x **a** crosses, FACS analysis revealed diploid DNA content in fusion products from XL280**a** *NAT* x XL280**a** *NEO*, *sxi2***a**Δ::*NAT* x *sxi2***a**Δ::*NEO*, and XL280**a** *NAT* x *sxi2***a**Δ::*NEO*, whereas haploid DNA content was found in the *sxi2***a**Δ::*NAT* x XL280**a** *NEO* cross (Fig 4D). To further confirm whether the donor cell directly participated in fusion events, genotyping PCR was performed to amplify *STE20***a** and *STE20*α, allelic variants of the same mating-type-specific genes located with the *MAT* locus. None of the independent fusion products contained the *STE20* allele from the pheromone donor’s mating type, indicating that any triploid products involving the donor strain are rare and were not observed in our analysis (Fig 4D and 4E).

To dissect the transcriptional differences underlying increased cell fusion frequency, we performed 3’ enriched RNA-sequencing for wildtype and *sxi1*αΔ bilateral Ménage à trois matings conducted on V8 (pH=7) for 72 hours, the identical conditions used for fusion frequency assays. Wildtype samples consisted of XL280α NAT x XL280α NEO x JEC20, while *sxi1*αΔ bilateral samples consisted of *sxi1*αΔ::*NAT* x *sxi1*αΔ::*NEO* x JEC20. Libraries were prepared with a 3’ enriched RNA-seq protocol incorporating unique molecular identifiers to enable deduplication and accurate transcript quantification, yielding an average of 11.75 million aligned reads per sample (S9 Fig). Principal component analysis revealed clear separation between wildtype and *sxi1*αΔ samples along PC1, which explained 63.2 percent of the variance, while PC2 accounted for 19 percent of the variance and showed partial overlap between groups (Fig 4F). Differential expression analysis with an adjusted p-value threshold of 0.05 or less (adjusted p-value<=0.05) and an absolute fold change cutoff of at least two (Fold Change >= 2) identified 134 differentially expressed genes, including 18 upregulated and 116 downregulated genes in *sxi1*αΔ bilateral matings relative to wildtype (Table S1). Consistent with previous findings, the known Sxi1α target *CLP1* was significantly downregulated in *sxi1*αΔ matings. Notably, the pheromone receptor gene *STE3*α and pheromone genes *MF*α*1–3* were significantly upregulated in *sxi1*αΔ matings (Fig 4G). Because genome-wide DEG analysis applies stringent statistical thresholds that may exclude biologically relevant mating and fusion genes, we next examined normalized expression levels using log2 counts per million (CPM) to directly compare transcript abundance across conditions. Examination of log2 CPM for mating and cell fusion related genes confirmed elevated expression of *MF*α*1–3* and *STE3*α, but not for the key transcription factor gene *STE12*α, in *sxi1*αΔ matings. Pheromone genes at the *MAT***a** locus, *MF***a***1**–**3*, were also upregulated, whereas *STE3***a** and *STE12***a** showed no significant changes (Fig 4H). In addition, significant expression differences were observed for additional Sxi1α targets such as *CPR2* and for key mating regulators *ZNF2* [51] and *CFS1* [52], while no significant differences were detected for *SXI2***a,** *MAT2*, *CVA1* and the fusion associated genes *PRM1* and *KAR5* [53] (Fig 4H).

### Sxi1α and Sxi2a regulate expression of unique gene subsets in unisexual crosses

To explore potential transcriptional regulatory roles for Sxi1α and Sxi2**a** in unisexual and α x **a** sexual reproduction, we performed RNA-seq of solo-cultures (XL280α, XL280**a**, *sxi1*αΔ, and *sxi2***a**Δ), and co-cultures including a wildtype cross (XL280α x XL280**a**), unilateral crosses (*sxi1*αΔ x XL280**a** and XL280α x *sxi2***a**Δ), and a bilateral mutant cross (*sxi1*αΔ x *sxi2***a**Δ). Reads were mapped to a concatenated assembly consisting of a chromosome-level XL280α genome and the XL280**a** *MAT***a** locus [54], with *MAT***a**- or *MAT*α-masked versions used for solo-cultures and the unmasked assembly for co-cultures. We acquired a mean of 22.4 million reads per sample (SD = 7.6 M) and achieved a mean alignment rate of 95.2% (SD = 0.8%). Genes with at least 1 transcript per million (TPM) in at least one sample were retained for downstream analysis. In both α and **a** solo-cultures, principal component analysis (PCA) of the 500 most variable transcripts showed that the presence/absence of *SXI1*α or *SXI2***a** explained the majority of the variance (S10A and S10B Fig). One replicate from the *sxi1*αΔ solo culture clustered separately from the others in both principal component space and in hierarchical clustering of the relative (scaled variance stabilizing transformation (vst) normalized) expression of the top 500 most variable genes (S10C Fig). Therefore, this replicate was excluded from downstream expression analysis. Re-analysis without this replicate resulted in reduced variance between replicates and a larger proportion of variance explained by PC1 (S10D Fig). All four co-culture crosses showed clear clustering of technical replicates and separation by genotype, indicating that transcriptional differences in these experiments are explained primarily by the independent variable with minimal contributions from confounding variables (S1E Fig). PC1 (62% variance) appears to be explained by the presence of a functional copy of both *SXI1*α and *SXI2***a**, with the samples from the wildtype cross clustering separately from the crosses where one or both genes were deleted. Interestingly, PC2 (25%) shows strong, discrete clustering by genotype, indicating transcriptional differences between crosses lacking *SXI1*α, *SXI2***a**, and both *SXI1*α and *SXI2***a** (S10E Fig).

We next performed differential gene expression analysis, treating genes with a false discovery rate (FDR)-adjusted p-value less than or equal to 0.05 and at least a twofold positive or negative expression change as significantly differentially expressed. In the *sxi1*αΔ comparison, *SXI1*α was identified as a DEG, and further analysis of the mapped RNA-seq reads showed that reads mapping to *SXI1*α were concentrated at the same regions with a high mapping rate of the Illumina DNA-seq reads (S11 Fig and Fig 2B). No reads were detected in the open reading frame. This suggests that the 5’ UTR region used for homologous recombination may have integrated ectopically in the genome and/or increased in copy number. A *MAT*α-adjacent gene, *FAO1*, was also identified as a DEG in the *sxi1*αΔ solo culture. Reads mapping to this gene were concentrated at the 3’ end that was also part of the homology region used for integration of the selectable marker during *SXI1*α gene deletion. It is therefore likely that this effect is also an artifact of ectopic integration of the resistance cassette. Based on these observations, *SXI1*α, *FAO1*, and *SXI2***a** were excluded from consideration as DEGs.

In solo-cultures, *sxi1*αΔ had 19 DEGs compared to XL280α, while *sxi2***a**Δ had 23 DEGs relative to XL280**a** (Fig 5A, 5B and Table S2). There were notably no shared DEGs between *sxi1*αΔ and *sxi2***a**Δ (Fig 5C). Amt2 (CNJ01880) [55], homologous to a well-characterized ammonium permease required for mating under low-nitrogen conditions in *C. neoformans*, was upregulated in the *sxi1*αΔ mutant. Gene ontology (GO) enrichment analysis of the identified DEGs in the *sxi1*αΔ mutant revealed two significantly enriched terms, *nucleobase transport* and *transmembrane transport* (S12A Fig), whereas no significant enrichment was observed for DEGs from the *sxi2***a**Δ mutant.

**Figure 5.**
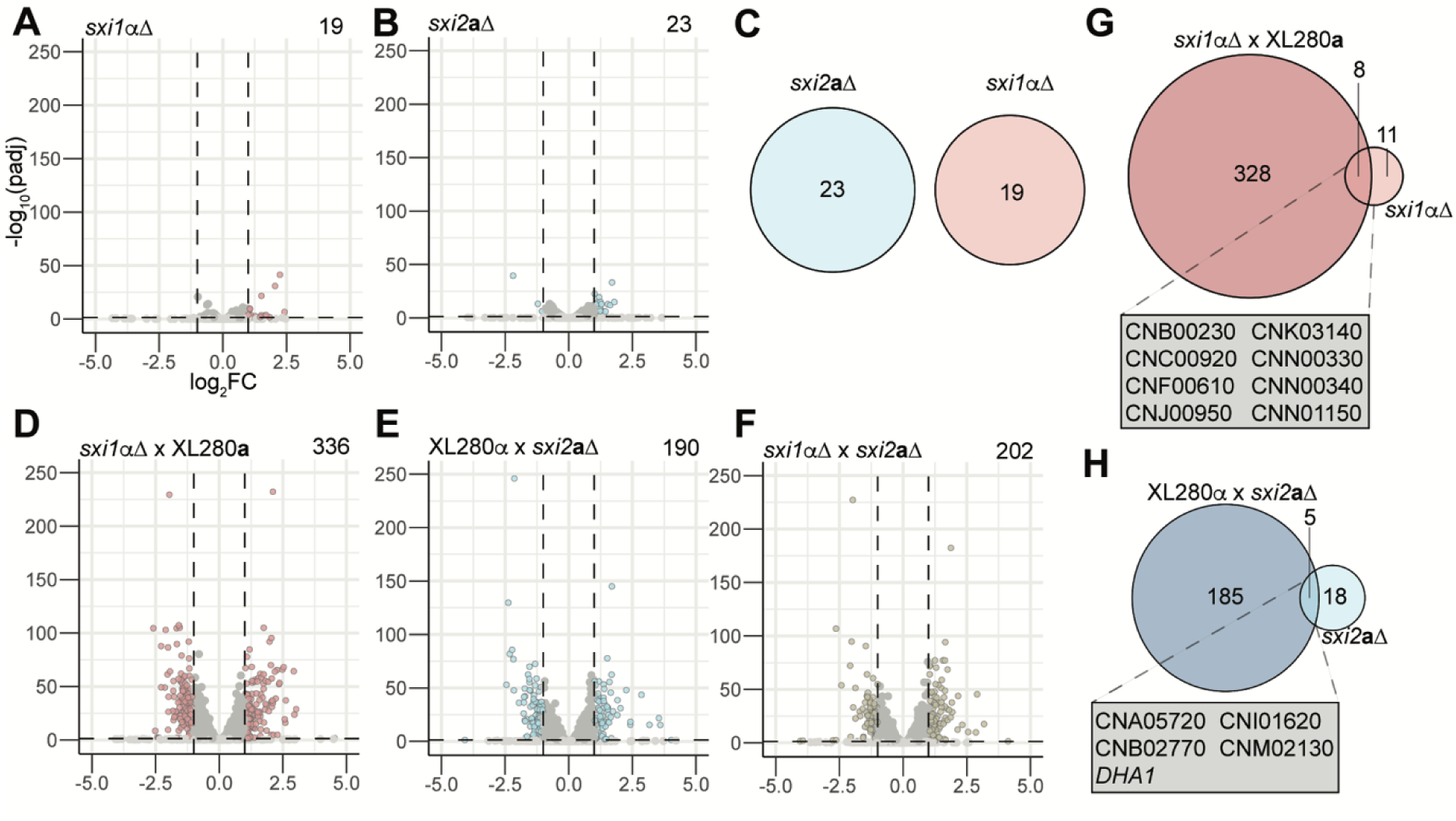
Sxi1α and Sxi2a regulate transcription of distinct subsets of genes during unisexual and bisexual reproduction. (A and B) Differentially expressed genes in *sxi1*αΔ (A) and *sxi2***a**Δ (B) relative to wild-type XL280α or XL280**a**, respectively, during solo cultures (72h at RT on V8 medium). Dashed lines indicate p_adj_ = 0.05 and log_2_FC = ± 1. (C) Comparison of DEGs in (A) and (B). (D-F) DEGs relative to XL280α x XL280**a** in crosses between *sxi1*αΔ x XL280**a** (D), XL280α x *sxi2***a**Δ (E), and *sxi1*αΔ x *sxi2***a**Δ (F). (G and H) Comparisons of DEGs in *sxi1*αΔ (G) and *sxi2***a**Δ (H) mutants between solo cultures and α x **a** crosses, with overlapping genes highlighted in the boxes.

### *SXI1*α and *SXI2*a have distinct and overlapping transcriptional effects in unisexual and α x a sexual crosses

Given the apparent α- and **a**- specific transcriptional roles of *SXI1*α and *SXI2***a** during unisexual reproduction, we next questioned whether these transcription factors may also function independently of one another, in addition to their known function as a heterodimer, during α x **a** sexual reproduction. We compared transcriptional changes in α x **a** crosses of opposite mating type XL280 parents deficient for one (unilateral cross) or both (bilateral cross) transcription factors. Comparing all mutant crosses to the wildtype cross XL280α x XL280**a**, we identified 336, 190, and 202 DEGs in *sxi1*αΔ x XL280**a**, XL280α x *sxi2***a**Δ, and *sxi1*αΔ x *sxi2***a**Δ, respectively (Table S2). In contrast to the solo-cultures undergoing unisexual reproduction, where the sets of DEGs were markedly smaller and primarily upregulated in *sxi1*αΔ and *sxi2***a**Δ relative to wildtype, DEGs had both increased and decreased expression relative to the XL280α x XL280**a** co-culture cross (Fig 5D-5F). GO term enrichment analysis revealed that DEGs from the unilateral cross *sxi1*αΔ x XL280**a** were predominantly associated with carbohydrate transport and metabolism, including monosaccharide, hexose, and glucose transmembrane transport, as well as carbohydrate catabolic processes (S12B Fig). The unilateral cross XL280α x *sxi2***a**Δ showed enrichment for broader carbohydrate metabolic and catabolic processes (S12B Fig). In the bilateral mutant cross *sxi1*αΔ x *sxi2***a**Δ, enriched terms were similar to those in the *sxi1*αΔ unilateral cross, with strong representation of carbohydrate transport, catabolism, and glucose-related processes (S12B Fig). These results suggest that Sxi1α and Sxi2**a** are involved in broad metabolic regulation during sexual reproduction, evidencing their functionality beyond processes directly implicated in sexual development. Further, observations that deletion of *SXI1*α has a more pronounced impact on carbohydrate-related pathways during mating compared to deletion of *SXI2***a** provide additional support for the existence of distinct roles for these transcription factors.

There were eight DEGs in common between solo *sxi1*αΔ cultures and the *sxi1*αΔ x XL280**a** cross and five DEGs in common between solo *sxi2***a**Δ and the XL280α x *sxi2***a**Δ cross (Fig 5G and 5H). These genes are largely uncharacterized, with functions that remain unknown, except for *DHA1* (CND04870) [56], a secreted protein whose expression is regulated by copper availability and which was identified as a DEG in both modes of sexual reproduction in the *sxi2***a**Δ mutant. Comparison of the predicted protein sequences of these shared genes with *S. cerevisiae* proteins shows potential orthologs with roles in metabolic processes (Table S3).

Given the enrichment for metabolic processes in the transcriptomic data, we used Biolog YT phenotypic plates to assess whether these changes translated into altered substrate utilization [57,58]. The Biolog YT panel is specifically designed for yeast metabolic phenotyping and contains 94 wells, each preloaded with a different carbon source, enabling comprehensive assessment of both respiratory and growth-based metabolic activity over time. PCA of metabolic activity at 72, 96, and 120 h showed partial separation between *sxi1*αΔ and XL280α (PC1 = 24.4% variance) (S13A Fig), with PERMANOVA confirming a significant difference (R² = 0.123, p = 0.026). *sxi2***a**Δ and XL280**a** showed greater overlap (PC1 = 43%) (S13B Fig), with no significant difference (R² = 0.057, p = 0.389). To pinpoint the substrates contributing to these differences, pairwise t-tests were performed for each carbon source at each time point. In the *sxi1*αΔ mutant, significant differences (p < 0.05) compared to XL280α were observed for utilization of D-gluconic acid, D-sorbitol, L-malic acid, N-acetyl-L-glutamic acid plus D-xylose, and palatinose at one or more tested time points (S14 Fig). In the *sxi2***a**Δ mutant, utilization of β-methyl-D-glucoside, gentibiose, quinic acid plus D-xylose, D-galactose, D-raffinose, D-sorbitol, and dextrin differed significantly from XL280**a** in at least one of the tested time points (S15 Fig). These phenotypic differences in carbon source utilization are consistent with the RNA-seq GO term enrichment results, which indicated that the Sxi1α and Sxi2**a** mutations affect genes involved in carbohydrate transport and metabolism.

The identification of shared DEGs between unisexual and α x **a** sexual crosses indicates that Sxi1α, and potentially Sxi2**a**, could have distinct transcriptional effects from their heterodimer interacting partner. These observations, coupled with the computational prediction and yeast two-hybrid assays that Sxi2**a** may be capable of forming homodimers, led us to question the extent of Sxi1α- and Sxi2**a**-dependent transcriptional effects occurring independently of the presence of a functional heterodimer interacting partner during α x **a** crosses. To differentiate between transcriptional events dependent on the Sxi1α-Sxi2**a** heterodimer as opposed to only Sxi1α or only Sxi2**a**, we examined the direction of differential expression in DEGs identified in one or more crosses (Fig 6A). Compared to the XL280α x XL280**a** cross, candidate heterodimer-regulated genes should undergo either a positive (heterodimer-repressed) or negative (heterodimer-activated) change in expression in crosses lacking *SXI1*α*, SXI2***a**, or both (Fig 6A, top panel).

**Figure 6.**
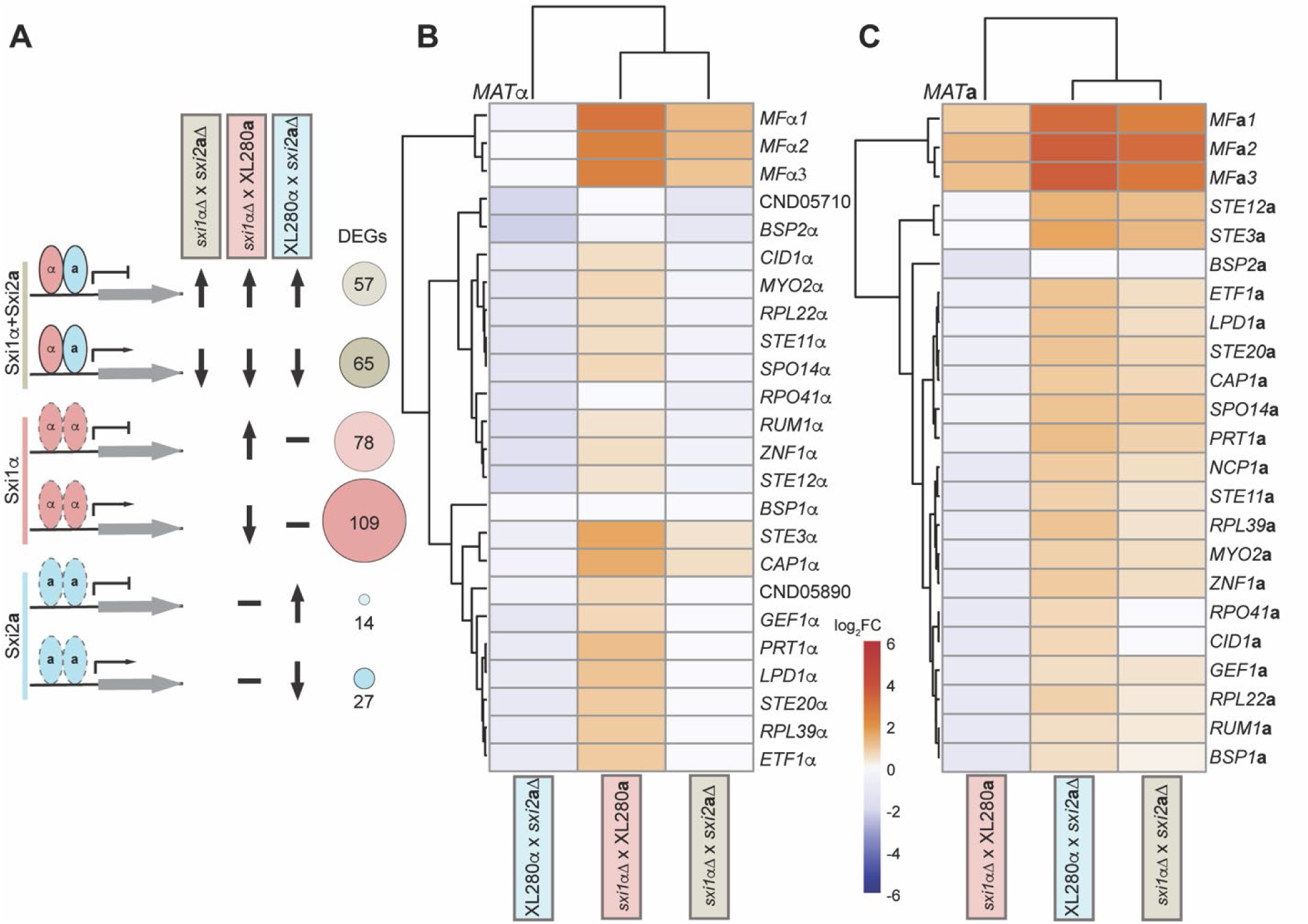
Sxi1α and Sxi2a control mating genes in distinct and opposing manners. (A) Schematic of 3 hypotheses for Sxi1α and Sxi2**a** function during α x **a** sexual reproduction. Genes in the first scenario are either positively or negatively regulated by the Sxi1α-Sxi2**a** heterodimer and undergo similar expression changes in all crosses lacking functional Sxi1α and/or Sxi2**a**. Genes in the second scenario are regulated by Sxi1α and are differentially expressed in crosses lacking a functional copy of Sxi1α but not in the XL280α x *sxi2***a**Δ cross because expression changes are independent of Sxi2**a**. Genes in the third scenario are differentially expressed in crosses lacking functional Sxi2**a** (XL280α x *sxi2***a**Δ and *sxi1*αΔ x *sxi2***a**Δ) but unchanged in the presence of a functional copy of Sxi2**a** (*sxi1*αΔ x XL280**a**). (B and C) log_2_FC expression values of *MAT*α (B) and *MAT***a** (C) genes in *sxi1*αΔ x XL280**a**, XL280α x *sxi2***a**Δ, and *sxi1*αΔ x *sxi2***a**Δ mutant crosses relative to XL280α x XL280**a** wildtype cross.

By these criteria, we identified 57 and 65 potential heterodimer-repressed and -activated genes, respectively (Fig 6A, top panel). Consistent with previous reports [26,29], *CLP1* (CNB01190) was among the set of heterodimer-activated genes. We next reasoned that any genes whose expression depends only on Sxi1α would have either increased (Sxi1α-repressed) or decreased (Sxi1α-activated) expression relative to wildtype in the *sxi1*αΔ x XL280**a** cross, but no expression change in the XL280α x *sxi2***a**Δ cross because XL280α has a functional copy of Sxi1α (Fig 6A, middle panel). Strikingly, we identified a larger gene set that fit these criteria than potential heterodimer-regulated genes, with 78 and 109 Sxi1α-activated and - repressed DEGs, respectively (Fig 6A, middle panel). Lastly, we applied similar reasoning to identify DEGs whose expression depends only on a functional copy of *SXI2***a** (Fig 6A, bottom panel). From this comparison, we identified 14 and 27 potential Sxi2**a**-activated and -repressed genes, respectively (Fig 6A, bottom panel). Surprisingly, *CPR2* (CNB04250) [27], a gene thought to be activated by the Sxi1α-Sxi2**a** heterodimer, was present in the set of 27 genes with potential Sxi2**a**-dependent activation.

### *SXI1*α and *SXI2*a have distinct regulatory effects on the α and a *MAT* loci

We observed an enrichment of *MAT*α and *MAT***a** genes in the Sxi1α- and Sxi2**a**-only dependent sets of DEGs compared to the heterodimer set, in which only two genes (*MF***a***1* and *MF***a***2*) were predicted to be heterodimer repressed (Fig 6A, top panel). Plotting expression change in all crosses for the *MAT*α and *MAT***a** loci revealed an even more surprising pattern, where a majority of genes in each locus appear to be controlled by Sxi1α and Sxi2**a** in opposing directions. *MAT*α genes tend to have increased expression in the absence of Sxi1α and decreased expression in the absence of Sxi2**a**, suggesting a negative regulatory role for Sxi1α and a positive regulatory role for Sxi2**a** (Fig. 6B). Conversely, *MAT***a** genes generally have positive expression changes in the absence of Sxi2**a** and negative expression changes in the absence of Sxi1α (Fig. 6C). In the case of *MAT***a** genes, expression differences in the *sxi1*αΔ x *sxi2***a**Δ crosses more closely mirror those in the XL280α x *sxi2***a**Δ than in *sxi1*αΔ x XL280**a**. Therefore, while *sxi2***a**Δ may result in a smaller set of uniquely differentially expressed genes, Sxi2**a** may have an important role in driving *MAT***a** gene expression change in opposition to Sxi1α. Taken together, these results support distinct roles for *SXI1*α and *SXI2***a** in both mating-type-specific and more general gene regulation.

## Discussion

The AlphaFold3-predicted heterodimer structure of Sxi1α and Sxi2**a** closely resembles the yeast **a**1-α2 complex [40] and the predicted *U. maydis* bW-bE heterodimer, consistent with prior studies [2,22]. This apparent conservation across diverse yeast species supports an evolutionarily important role of the Sxi1α-Sxi2**a** complex in α x **a** mating. Lower-confidence predictions indicate that Sxi1α and Sxi2**a** may also form homodimers, similar to α2 in *S. cerevisiae*, raising the possibility of independent, non-heterodimer functions. Importantly, results from yeast two-hybrid assay support functional asymmetry between the two HD proteins. Whereas HD2 Sxi2**a** appears capable of forming a homodimer, comparable homodimerization by HD1 Sxi1α has not been observed. This distinction suggests that Sxi2**a** may possess a greater capacity for autonomous regulatory activity, while Sxi1α may depend more strongly on alternative co-factors or context-specific interactions. Such asymmetry provides a potential mechanistic explanation for the opposing and gene-specific regulatory effects of Sxi1α and Sxi2**a** observed across multiple mating contexts. Interestingly, homodimerization of HD2 has been reported in the homothallic fungus *Cystofilobasidium capitatum*, where HD2 homodimers can substitute for the canonical HD1-HD2 heterodimer and thereby provide an alternative mechanism for regulating sexual reproduction [59]. Likewise, in the basidiomycetous yeast *Phaffia rhodozyma*, the homeodomain proteins HD1 and HD2 display only a weak heterodimer interaction, while experimental assays revealed potential homodimer formation [60]. In contrast, in *Ustilago maydis* the bE and bW homeodomain transcription factors strictly function through non-self heterodimerization, and no evidence for functional homodimerization has been found [61]. These examples suggest that HD homodimerization, although rare, may provide lineage-specific solutions for regulating sexual development. Together with our transcriptomic findings, this led us to hypothesize that these transcription factors can act both together and separately, whether as homodimers or by another means regulating mating before and after cell-cell fusion in unisexual and α x **a** reproduction.

Our transcriptomic results provide further evidence for this model, as we observed Sxi2**a**-independent transcriptional regulation by Sxi1α and Sxi1α-independent regulation by Sxi2**a** in both unisexual and α x **a** sexual matings. In addition to regulating largely distinct gene sets, Sxi1α and Sxi2**a** exert contrasting effects on the two mating-type loci. We observed an enrichment of *MAT*α and *MAT***a** genes in the Sxi1α-only and Sxi2**a**-only dependent DEG sets in α x **a** crosses, whereas the heterodimer-dependent set contained only two repressed genes (*MF***a***1* and *MF***a***2*). Strikingly, most *MAT*α genes showed increased expression in the absence of *Sxi1*α but decreased expression in the absence of *Sxi2***a**, suggesting that Sxi1α functions primarily as a negative regulator and Sxi2**a** as a positive regulator at this locus. The pattern was reversed for *MAT***a** genes, with *Sxi1*α acting positively and *Sxi2***a** negatively. These results indicate that, beyond their shared role as a heterodimer, Sxi1α and Sxi2**a** also engage in mating-type-specific regulatory programs that act in opposition to each other, potentially fine-tuning the expression of *MAT*-linked genes during α x **a** sexual reproduction. Deletion of Sxi1α caused an increase in α-α cell fusion, consistent with a functional role beyond that of Sxi1α-Sxi2**a** heterodimer. To further interpret the molecular basis of altered fusion frequency, we considered transcriptional changes associated specifically with fusion-competent mating conditions. Under these conditions, genes involved in pheromone sensing and signaling, including the pheromone receptor gene *STE3* and multiple pheromone genes, were upregulated, whereas canonical membrane fusion and karyogamy genes such as *PRM1* and *KAR5* [53] did not show corresponding induction. This pattern suggests that enhanced fusion may arise from elevated mating competence or increased responsiveness to pheromone cues rather than from direct transcriptional activation of the fusion machinery itself. Such regulation at the level of upstream signaling could provide a flexible mechanism for modulating fusion probability without broadly altering the core fusion apparatus. These results support the possibility that Sxi1α may act independently of Sxi2**a**, and future work will continue to explore potential co-factors/interacting partner(s). In α x **a** reproduction, both Sxi1α and Sxi2**a** are essential for normal hyphal development and sporulation in multiple genetic backgrounds [2,22]. In contrast, in the XL280 background, deletion of either gene did not impair unisexual colony morphology or sporulation. These results, coupled with the observation of *SXI1*α - and *SXI2***a**-dependent transcriptional activity during unisexual and α x **a** sexual reproduction, suggest that these transcription factors are active but may play distinct roles in α x α and **a** x **a** versus α x **a** reproduction, and that unisexual reproduction is likely regulated in a manner that is at least partially distinct from α x **a** sexual reproduction. This is also consistent with recent reports that Sxi2**a** is dispensable for unisexual reproduction in *C. neoformans* [31].

Functional differences between the two HD proteins were most evident in unisexual fusion assays. Sxi1α inhibited fusion in both unilateral and bilateral mutant crosses, whereas Sxi2**a** inhibition was observed only in unilateral crosses. This may indicate that Sxi2**a** plays a less-prominent or context-dependent role in **a** x **a** cell fusion, possibly masked in bilateral crosses by redundancy with other transcription factors. Testing this possibility will require identifying Sxi2**a**’s non-Sxi1α partners and determining their contributions to fusion regulation.

Our transcriptomic analysis identified distinct subsets of Sxi1α- and Sxi2**a**-regulated genes in both unilateral and bilateral crosses. A majority of these genes remain uncharacterized, inviting future functional investigation into their potential involvement in cell-cell fusion and sexual reproduction. Notably, GO term enrichment highlighted metabolism-related processes, and Biolog YT plate profiling revealed subtle but reproducible shifts in carbon source utilization between wildtype and mutants. Together, these findings suggest that Sxi1α and Sxi2**a** influence metabolic pathways, which may, in turn, modulate mating efficiency under nutrient-variable conditions, thereby linking metabolic state to the mating regulatory network and highlighting carbohydrate metabolism as an important area for future investigation of Sxi1α and Sxi2**a** function.

By analyzing 341 *MAT*α clinical and environmental isolates out of the total 387 isolates in the *C. neoformans* Strain Diversity Collection [33] through a similar pipeline [62] as reported in our previous work [63], we found there were only two closely related environmental isolates (Muc504-1 and Muc489-1) that harbored an identical high-impact nonsense mutation in the *SXI1*α gene, resulting in truncation of the C-terminal 23 amino acids. The widespread conservation of this gene suggests that it is under positive selection and likely has important functions in a variety of ecological contexts. Unisexual reproduction, which allows cells to produce stress-resistant spores under harsh environmental conditions in the absence of an opposite mating type partner, provides one hypothetical avenue by which sex-specific genes like *SXI1*α could continue to be selected for in a predominantly *MAT*α population. While the inhibitory role of Sxi1α on unisexual α-α fusion could, in principle, be expected to limit the expansion of *MAT*α lineages and favor *MAT***a** cells, this prediction appears inconsistent with the observed predominance of *MAT*α in natural populations. This paradox suggests that additional mechanisms beyond Sxi1α regulation must contribute to shaping the observed mating-type distribution. Further studies are needed to clarify the downstream regulators involved in the novel repressive role of Sxi1α in unisexual reproduction and other potential mating-independent functions. These lines of investigation have the potential to elucidate fitness-enhancing strategies employed by this organism to undergo unisexual meiosis and produce infectious spores. Over evolutionary time, it is conceivable that selection for alternative reproductive strategies such as unisexual reproduction could have contributed to the widespread dominance of *MAT*α strains. In this context, we speculate that repression of α–α cell fusion by the α-specific homeodomain protein Sxi1α represents a *Cryptococcus*-specific evolutionary adaptation that fine-tunes unisexual development to permit efficient spore production in the absence of a conidiospore-based asexual program. Such a mechanism may balance the selective advantage of abundant basidiospore formation with controlled mating behavior under partner-limited conditions.

In *S. cerevisiae*, the α2 transcription factor functions both as a homodimer and heterodimer with **a**1, playing key roles in mating-type regulation [6–9]. The transcriptional regulation of **a**-specific and haploid-specific genes involves interactions with the transcription factor Mcm1 [6,43]. Given this precedent, a similar dual functionality with/without mating partner may exist for the sex-specific HD transcription factor Sxi1α in *Cryptococcus*, and non-Sxi2**a** interacting partners may exist. Interestingly, orthologs of *S. cerevisiae* Mcm1 are also present in *C. neoformans* and *C. deneoformans*, where they have been reported to function in α x **a** sexual reproduction [49]. However, our functional analysis suggests that the *Cryptococcus* Mcm1 ortholog does not play a central role in *Cryptococcus* mating under the conditions examined. In contrast to its well-established function in budding yeast, loss of Mcm1 did not result in clear defects in unisexual or α x **a** sexual reproduction. Together with the absence of strong evidence for direct association with either Sxi1α or Sxi2**a**, these observations indicate that *Cryptococcus* mating circuits have diverged substantially from the canonical budding yeast α2–Mcm1 regulatory framework, relying instead on alternative or redundant transcriptional circuits. Our results suggest that Sxi1α might act as both a positive (e.g., in α x **a** mating development) and a negative regulator (e.g., in α x α cell fusion and over transcription of certain *MAT* genes) [23]. This phenomenon is reminiscent of the mitogen-activated protein kinase Kss1 in *S. cerevisiae*, which regulates pseudohyphal differentiation through complex signaling mechanisms and functions as a multifunctional regulator in fungal development with counter-balanced positive and negative roles [64]. Efforts to identify potential Sxi1α - and Sxi2**a** - specific transcription factor binding sites in the promoter regions of differentially expressed genes were not successful, likely due to the presence of numerous derivative consensus sequences (similar to what has previously been observed in *C. neoformans*), as well as the potential activation of regulatory cascades as a result of changes in Sxi1α and Sxi2**a** activity, meaning that many promoters of DEGs are not bound by the homeodomain proteins themselves. Future efforts to elucidate direct targets of the heterodimer and putative homodimers will involve ChIP-seq–based experimental validation of binding sites in the presence and absence of each transcription factor, ideally integrated with transcriptomic analyses to distinguish direct from indirect regulatory effects.

While the HD proteins Sxi1α and Sxi2**a** are dispensable for colony morphology, sporulation, and cell viability during unisexual reproduction, Sxi1α plays a key inhibitory role in α x α cell fusion, and the contribution of Sxi2**a** to this process appears to be more limited. RNA-seq analysis further revealed that the heterodimer partners also have distinct and independent regulatory programs, supporting functional roles beyond their shared activity as a complex. These findings suggest that Sxi1α may have novel functions during *Cryptococcus* mating that are yet to be characterized, potentially analogous to those in other well studied fungal species, and highlight an additional facet in the complex regulation of sexual reproduction in this important human fungal pathogen. Our findings also raise important questions regarding the functional conservation of sex-specific homeodomain proteins in ascomycetes and basidiomycetes. Future studies will investigate protein complexes involving Sxi1α, Sxi2**a**, and additional interacting partners to elucidate the extent of conservation with well-characterized homeodomain complexes such as **a**1-α2 in *S. cerevisiae*. Collectively, these findings support a model in which Sxi1α and Sxi2**a** act through partially independent regulatory programs that modulate pheromone signaling, transcriptional responsiveness, and developmental timing, rather than functioning exclusively as an obligate heterodimeric switch. Taken together, our findings demonstrate that gene regulation by Sxi1α and Sxi2**a** is more complex than previously appreciated and provide a foundation for future molecular analyses aimed at further delineating the *Cryptococcus* developmental program.

## Materials and methods

### Fungal strains and growth conditions

*C. deneoformans* and *C. neoformans* strains for this study are listed in Table S4. The strains were stored in 15% glycerol at -80°C for long-term storage. Fresh strains were streaked from glycerol stocks and maintained on solid YPD medium (1% yeast extract, 2% Bacto Peptone, 2% dextrose) at 30°C.

Spot dilution assays were performed with overnight cultures of the tested strains grown in liquid YPD at 30°C. Strains were diluted to an OD_600_ of 0.5 in sterile water. These cultures were further serially diluted 5-fold and spotted (3 μL for each dilution) onto YPD plates. The plates were incubated at 30°C and 37°C (testing thermotolerance) for 2 days before imaging.

### Protein structural predictions

Protein structure predictions were made with AlphaFold3 [65] on the AlphaFold server (https://alphafoldserver.com/). The predicted models and error plots were further processed with Chimera X (v 1.9) [66] for predicted local distance difference test (pLDDT) value filtering, alignment, and visualization.

### Yeast two-hybrid assay

Sxi1α, Sxi2**a**, and Sxi2**a**N constructs were generated previously [2]. Briefly, *SXI1*α and *SXI2***a** coding sequences from XL280 were cloned from cDNA into Gal4 DNA-binding domain (GBD) and Gal4 activation domain (GAD) vectors, and truncated *SXI2***a** fragments were generated by PCR amplification of the defined N-terminal region (encoding amino acids 1-217). For this study, the putative *C. deneoformans MCM1* coding sequence was amplified from XL280 cDNA and cloned into GBD and GAD vectors to generate yeast two-hybrid constructs. All previously generated and newly constructed plasmids were sequence-verified prior to use (Table S4). Reporter strains Y2HGold and Y187 were transformed with plasmids via the high-efficiency transformation method [67]. Diploids were generated by mating and selected for on synthetic dextrose (SD) dropout medium (0.67% yeast nitrogen base, 2% dextrose) minus leucine and tryptophan. Adenine sulfate was added to the medium at a concentration of 20 mg/L, and amino acids and uracil were included at standard concentrations to support auxotrophic growth requirements. Gal4-dependent activation of *ADE2* and *HIS3* reporter genes was assessed by growth on SD medium additionally lacking adenine or histidine, respectively. β-galactosidase activity was measured using a chlorophenol red-β-D-galactopyranoside (CPRG)–based liquid assay (OZ Biosciences) according to the manufacturer’s instructions. Absorbance was measured at 595 nm, and enzyme activity (milliunits) was calculated based on a linear standard curve.

### Gene deletion and confirmation

Approximately 1.0 kb 5’ and 3’ flanking regions of the *SXI1*α gene and 60 bp 5’ and 3’ flanking regions of the *SXI2***a** and *MCM1* gene were constructed with primers listed in Table S5 to target their respective genes in XL280 for homologous recombination [46]. Overlap PCR with a selectable marker was used to construct the DNA donor with 1.0 kb homology, while 60 bp microhomology was included in the primers for donor amplification. *C. neoformans* codon optimized Cas9, gRNAs (a single gRNA targeting *SXI1*α and two gRNAs targeting *SXI2***a**), and drug resistance cassettes (*NAT* from pAI3 or *NEO* from pJAF1) containing homologous flanking sequences were introduced into XL280 wildtype strains via the previously reported Transient CRISPR-Cas9 Coupled with Electroporation (TRACE) system [38,39]. Transformants were screened with PCR internal to the ORF, and 5′ and 3′ junction PCR to the drug resistance marker integration sites to confirm integration at the genomic locus. Illumina whole-genome sequencing was performed on positive transformants with Novaseq X Plus platform (2x150 bp). The sequencing reads were mapped to a modified version of the XL280α genome [54] with the Burrows-Wheeler Alignment tool (BWA) (v 0.7.17) [68] to further confirm the deletion mutant strains. The modification included adding the *MAT***a** locus sequence of JEC20 (GenBank: AF542530.2) as an additional contig.

H99 *mcm1*Δ was obtained from [49] and crossed with KN99**a** to obtain KN99**a** *Mcm1*Δ. Both mutants were verified by internal PCR targeting the *MCM1* ORF with the primers listed in Table S5.

### Capsule formation and melanin production assays

Tested strains, with an initial OD_600_ of 0.1 in 4 mL liquid RPMI medium (RPMI 1640 with 2% dextrose), were incubated on a roller drum at 70 rpm and 30°C for 3 days. After washing and resuspension in water, the cells were negatively stained with an equal volume of India ink to visualize the capsules.

To assess melanin production, overnight saturated cultures of test strains were spotted (10 μL) onto Niger seed agar plates (7% Niger seed, 0.1% dextrose). The Niger seed plates were incubated at 30°C and 37°C for 8 days before photographing.

### Competition assays

Overnight cultures of tested strains grown in liquid YPD at 30°C were diluted to OD_600_ of 0.2 and equally mixed in a 1:1 ratio in 5 mL of liquid YPD for co-culture. *sxi1*αΔ::*NAT* mutants (resistant to NAT) and *sxi2***a**Δ::*NEO* mutants (resistant to NEO) were mixed with XL280α and XL280**a**, respectively. The mixture was plated on YPD plates, YPD + nourseothricin (NAT), and YPD + neomycin (NEO) to confirm starting ratio. After 24 and 48 hours of co-culture, an aliquot of the mixture was sampled, and plating was repeated with OD_600_ diluted to 0.1 to track changes in relative abundance of mutants. Four independent replicates were performed for each competition.

For the competition assay performed on MS medium, overnight cultures were similarly diluted to OD_600_ of 1. *sxi1*αΔ::*NAT* or *sxi2***a**Δ::*NEO* mutants and the corresponding XL280 wildtype strain were equally mixed in a 1:1 ratio and spotted on MS medium. Spores were dissected 10 to 14 days post spotting. Yeast cells from internal regions were also collected. Spores or yeast cells were further transferred to YPD+NAT, YPD+NEO and YPD to determine the ratio of mutants in the population. Five independent spots were investigated for each competition.

### Mating and cell-cell fusion assay

Mating plates were set up as described previously [69]. α x α and **a** x **a** unisexual reproduction assays were performed on Murashige and Skoog (MS, Sigma, USA) plates for 1 to 2 weeks before imaging and dissection. Basidiospores generated during unisexual reproduction were randomly dissected and plated on YPD plates for 2 to 3 days to determine spore viability and germination rates.

Cell-cell fusion frequency during α x α unisexual outcrossing in XL280α *NAT* (XL561 and XL562) and XL280α *NEO* (XL563 and XL564) was compared to bilateral mutant crosses of *sxi1*αΔ::*NAT* and *sxi1*αΔ: *NEO* or unilateral crosses of *sxi1*αΔ::*NAT* and XL280α *NEO*. Similarly, **a** x **a** unisexual outcrossing in XL280**a** *NAT* (CF1816) and XL280**a** *NEO* (JHG223) was compared to bilateral mutant crosses of *sxi2***a**Δ::*NAT* and *sxi2***a**Δ::*NEO* or unilateral crosses of *sxi2***a**Δ::*NAT* and XL280**a** *NEO* or *sxi2**a***Δ::*NEO* and XL280**a** *NAT*.

Cells were diluted in sterile water to an OD_600_ of 1. Two *MAT*α strains (α x α outcrossing) or *MAT***a** strains (**a** x **a** outcrossing) containing either *NAT* or *NEO* resistance cassettes were mixed equally with JEC20 (*MAT***a**) or JEC21 (*MAT*α) for Ménage à trois matings on V8 agar (pH=7) and incubated for 72 hours in the dark at room temperature. The cells were then collected and plated on YPD + NAT (100 μg/mL) + NEO (200 μg/mL) and YPD alone to determine the ratio of colonies with dual resistance four to five days post-plating (i.e., fusion product) out of the total cells. Genotypes at mating type locus of the fusion products was validated via *STE20***a** or *STE20*α genotyping PCR.

### Flow cytometry analysis

Ploidy of the fusion products from α x α or **a** x **a** cell fusion was determined by fluorescence-activated cell sorting (FACS) analysis as previously described [70]. Cells were grown overnight on solid YPD medium at 30 °C, harvested, and washed with 1 mL PBS. The cells were fixed in 70% ethanol at 4 °C overnight, pelleted and washed with 1 x NS buffer (10 mM Tris-HCl pH=7.5, 0.25 M sucrose, 1 mM EDTA, 1 mM MgCl_2_, 0.13 mM CaCl_2_, 0.1 mM ZnCl_2_, 0.5 mM phenylmethylsulfonyl fluoride, 7 mM β-mercaptoethanol). Samples were treated with RNase A (1.05 mg/mL) and stained with propidium iodide (PI, 30 μg/mL) in a total volume of 200 μL 1x NS solution. Stained cells were diluted in 1x Tris-PI buffer (0.964 M Tris-HCl pH=7.5, 35 µg/mL PI) prior to analysis. FACS was performed at Duke Cancer Institute Flow Cytometry Core Facility with approximately 10,000 cells analyzed per sample. Flow cytometry data were visualized using FlowJo (v.10.10.0).

### Light and scanning electron microscopy (SEM)

Mating structures from MS plates were imaged with an AxioSkop 2 fluorescence microscope to visualize hyphae, basidia and basidiospores under brightfield. SEM samples were prepared following a previously described protocol [50] with minor modifications. Briefly, a ∼2 cm x 2 cm agar plug containing patch of mated cells from MS plates was fixed in PBS containing 4% formaldehyde and 2% glutaraldehyde at 4°C for 2 hours. The fixed samples were then gradually dehydrated in a graded ethanol series (30%, 50%, 70%, 95%), with each step performed at 4°C for 15 min. This was followed by three washes in 100% ethanol, each at 4°C for 15 min. The samples were further dehydrated with a Ladd CPD3 Critical Point Dryer and subsequently coated with a thin layer of gold with a Denton Desk V Sputter Coater (Denton Vacuum, USA). Hyphae, basidia and basidiospores were observed with a scanning electron microscope equipped with an EDS detector (Apreo S, ThermoFisher, USA).

### RNA preparation and sequencing

For Ménage à trois matings, cells were diluted in sterile water to an OD_600_ of 1. Two *MAT*α strains (α x α outcrossing) containing either *NAT* or *NEO* resistance cassettes were mixed equally with JEC20 (*MAT***a**) on V8 agar (pH = 7) and incubated for 72 hours in the dark at room temperature after harvesting. Three biological replicates were performed for each strain or mating pair. Harvested samples were lyophilized and processed for RNA extraction using the RNeasy Mini Kit (Qiagen), following the manufacturer’s instructions. Library preparation and sequencing was performed by Plasmidsaurus (USA). RNA libraries were generated from total RNA using a strand-specific 3’ end counting protocol with unique molecular identifiers (UMIs) to enable accurate transcript abundance quantification (https://plasmidsaurus.com/rna). Libraries were sequenced on the Illumina platform, yielding deduplicated 3’ end enriched reads for gene expression analysis.

For both unisexual and α x **a** sexual crosses, strains were collected from 2 to 3-day-old cultures grown on solid YPD medium and diluted to an OD_600_ of 1. For unisexual crosses, 10 μL of the diluted culture (OD_600=_1) was spotted onto V8 agar plates (pH= 7), with 10 spots per plate and three technical replicate plates per biological replicate. For α x **a** crosses, equal volumes of *MAT*α and *MAT***a** cell suspensions (each at OD_600=_1) were mixed prior to spotting under the same conditions. Three biological replicates were performed for each strain or mating pair. Plates were incubated at room temperature for 72 hours in the dark, after which cells were washed off and collected. Harvested samples were lyophilized and processed for RNA extraction using the RNeasy Mini Kit (Qiagen), following the manufacturer’s instructions.

RNA was sequenced by the Duke University Sequencing and Genomic Technologies Core Facility. Poly-A enriched libraries were prepared from samples with RIN ≥ 7 using the Kapa mRNA-seq HyperPrep kit (Kapa Bioscience). Pooled libraries were sequenced on an Illumina Novaseq X Plus instrument in 150 bp paired-end reads.

### RNA-sequencing data analysis

For Plasmidsaurus RNA-seq, data was analyzed following Plasmidsaurus’s instructions. Briefly, raw reads were quality filtered by fastp (v0.24.0) [71], then aligned to the XL280α reference genome with an additional *MAT***a** sequences from JEC20 using STAR aligner (v.2.7.11b) [72], with non-canonical splice junction disallowed. PCR duplication was removed through UMI-based deduplication with UMIcollapse (v1.1.0) [73]. Alignment quality metrics, strand specificity, and read distribution across genomic features including enrichment toward transcript 3’ ends were assessed using RSeQC (v5.0.4) [74] and Qualimap (v2.3) [75], with results aggregated into a comprehensive quality control report using MultiQC (v1.32) [76]. Gene-level expression quantification was performed by using featureCounts (v2.1.1) [77]. PCA were calculated on normalized counts (TMM, trimmed mean of M-values) using Pearson correlation. Differential expression was done with edgeR [78] (v4.0.16) using standard practice including filtering for low-expressed genes with edgeR::filterByExprwith default values. Differentially expressed genes (DEGs) were identified using a significance threshold of adjusted p-value <= 0.05 and an absolute log_2_ fold change cutoff of 1. FeatureCounts gene level counts were filtered and TMM normalized using edgeR, and log_2_ counts per million (CPM) values were calculated. Significance was determined using Welch two sample t-tests with Benjamini Hochberg false discovery rate correction.

For RNA-seq data from both unisexual and α x **a** sexual crosses, Illumina universal adaptors were trimmed from paired reads with CutAdapt (v4.9) [79]. Trimmed reads were aligned to the XL280α + *MAT***a** concatenated reference assembly. Modified assemblies with *MAT*α (Chr4 coordinates 1522691-1656182) and *MAT***a** (MATa coordinates 0-148067) masked were generated by using the bedtools (v2.31.1) maskfasta function on the concatenated assembly [80]. XL280α background unisexual crosses were mapped to the *MAT***a**-masked assembly, XL280**a** background unisexual crosses to the *MAT*α-masked assembly, and α-**a** sexual crosses to the full, unmasked assembly with HISAT2 (v2.2.1) [81]. Counts per gene were quantified with featureCounts (v2.0.6) [77] with a gff annotation file generated by Liftoff (v1.6.3) [82] of annotated features from JEC20 and JEC21 [83].

Exploratory data analysis and differential expression analysis were performed with DESeq2 (v1.44.0) [84]. Prior to analysis, transcript counts were normalized by transcripts per million (TPM) within each replicate and genes with at least 1 TPM in 1 sample were retained. Clustering and principal component analyses were performed using vst normalized values for the top 500 most variable genes generated by DESeq2. Differential expression analysis was carried out in DESeq2 with alpha = 0.05 and a log_2_FC cutoff of +/- 1 was applied to identify DEGs from the resulting expression data. The R packages EnhancedVolcano (v1.22.0) (https://github.com/kevinblighe/EnhancedVolcano) and pheatmap (v1.0.13) (https://github.com/raivokolde/pheatmap) were used for data visualization. DEG sets were analyzed with the FungiDB (https://fungidb.org/fungidb/app) GO Enrichment tool using the JEC21 genome as background. Significance was assessed using Fisher’s exact test with Benjamini-Hochberg correction, retaining terms with FDR < 0.05. Enriched terms were visualized in RStudio as dot plots ranked by log₂(fold enrichment) or FDR, colored by −log_10_(FDR), and annotated with gene counts.

### Biolog phenotypic assays

Overnight cultures grown on solid YM agar (5 g peptone, 3 g yeast extract, 3 g malt extract, 10 g dextrose, 20g agar, pH=6.2) were collected and diluted in sterile water to an OD_600_ of 0.2. A volume of 100 μL of the diluted samples was inoculated into each well of the Biolog Yeast Identification Test Panel (YT Microplate) and cultured at 28 °C in the dark. Three biological replicates were performed for each strain.

Cell growth was monitored daily with SpectraMax iD3 plate reader at OD_590_ (oxidation test) and OD_600_ (Assimilation test). Growth on each substrate was normalized using the formula (Abs_sample - Abs_water)/(Abs_glucose-Abs_water). Principal component analysis (PCA) was performed using scaled data from Biolog YT plate assays. Differences in global metabolic profiles between genotypes were assessed using PERMANOVA (adonis2 function, vegan R package), with 999 permutations and Euclidean distance as the dissimilarity metric.

### Statistical analysis

Spore germination and unisexual competitiveness were analyzed with paired or unpaired student t-tests in Rstudio (version 2024.04.1) with the “t.test” function with argument “paired = TRUE” for paired tests or “paired = FALSE” for unpaired tests.

Fusion efficiency data were analyzed in RStudio (version 2024.04.1). Welch’s t-tests were performed using the t.test function with the argument “var.equal = FALSE” to compare fusion rates between selected sample groups. Mean values and standard deviations were plotted using ggplot2, with individual data points overlaid.

## Data availability

Raw WGS sequencing and RNA-seq reads have been deposited in Bioproject: PRJNA1219566. Scripts associated with RNA-seq data from both unisexual and α x **a** sexual crosses are available at: https://github.com/annalehmann/XL280_Sxi_transcriptome. Scripts related to Plasmidsaurus RNA-seq analysis are available at: https://github.com/Jhuang90/Huang-et-al.-Sxi1_Plasmidsaurus-RNA-seq-script.

## Acknowledgments

We thank Anna Floyd Averette for constant support, Connor Larmore for critical reading, Dr. Marco Dias Coelho for constructive criticism on the manuscript and suggestions on data presentation, Moayad Shehadeh and Dr. Zhuyun Bian for technical support, and all of the members of the Heitman Lab for constructive suggestions. We also thank Dr. Xiaorong Lin (University of Georgia) for sharing the XL280 congenic strains, Dr. Ruiyun Zeng (North Carolina State University) for assistance in data visualization, Dr. Paul Magwene and Claudia Zirión Martinez (Department of Biology, Duke) for sharing unpublished pipeline for variant detection from Strain Diversity Collection, Dr. Bin Li (Duke Cancer Institute Flow Cytometry Core Facility), Julian Liber (Department of Biology, Duke), and Dr. Devi Swain Lenz (Duke’s Sequencing and Genomic Technologies Core Facility) for advice and expertise.

## Funding

This study was supported by NIH/NIAID R01 grants (AI039115-28, AI050113-20, and AI133654-07. to J. Heitman) J. Heitman is co-director and fellow of the Canadian Institute for Advanced Research (CIFAR) program Fungal Kingdom: Threats & Opportunities. The funders had no role in study design, data collection and analysis, decision to publish, or preparation of the manuscript.

## Competing interests

I have read the journal’s policy and the authors of this manuscript have the following competing interests:. All other authors have declared that no competing interests exist.

## Supporting information captions

Table S1 DEGs identified during Ménage à trois matings.

Table S2 DEGs identified under different mating comparisons.

Table S3 DEGs shard between unisexual and α x **a** sexual reproduction.

Table S4 Strains and plasmids used in this study.

Table S5 Primers used in this study.

## Notes

### Competing Interest Statement

The authors have declared no competing interest.

### Summary of Updates

We have carefully and thoroughly addressed all comments and incorporated the valuable suggestions. This revision includes several additional experiments and data analysis, including yeast two-hybrid assays, RNA-seq analysis, and the generation of new deletion mutants, which have substantially strengthened the manuscript.

