## Supplemental Figure for "Homeodomain protein Sxi1α independently controls cell-cell fusion and gene expression during sexual reproduction in *Cryptococcus deneoformans*"

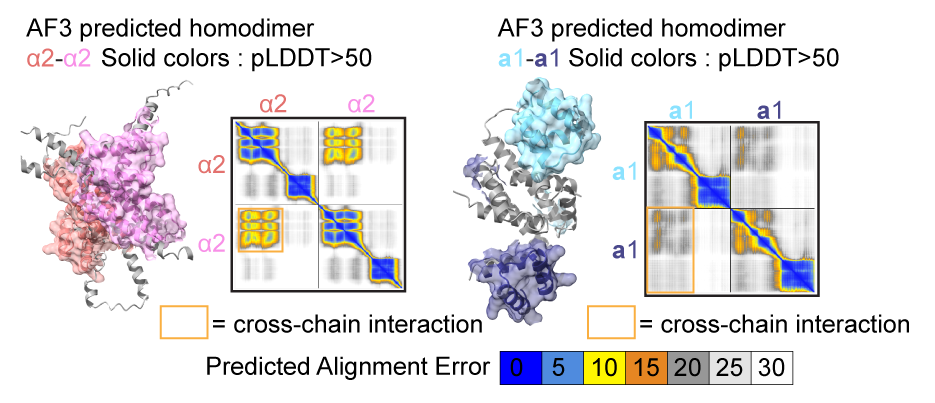

**Supplemental Figure 1. AlphaFold3 benchmarking of homodimer predictions using yeast HD regulators α2 and a1.**

Shown are predicted homodimer structures of *S. cerevisiae* α2 (left) and **a**1 (right) generated by AlphaFold3. Solid coloring indicates regions with pLDDT > 50. Predicted alignment error maps are shown with orange boxes highlighting cross chain interactions. A well-defined low-PAE interface is observed for α2, whereas **a**1 shows weak or poorly defined cross-chain interactions, consistent with experimental evidence.

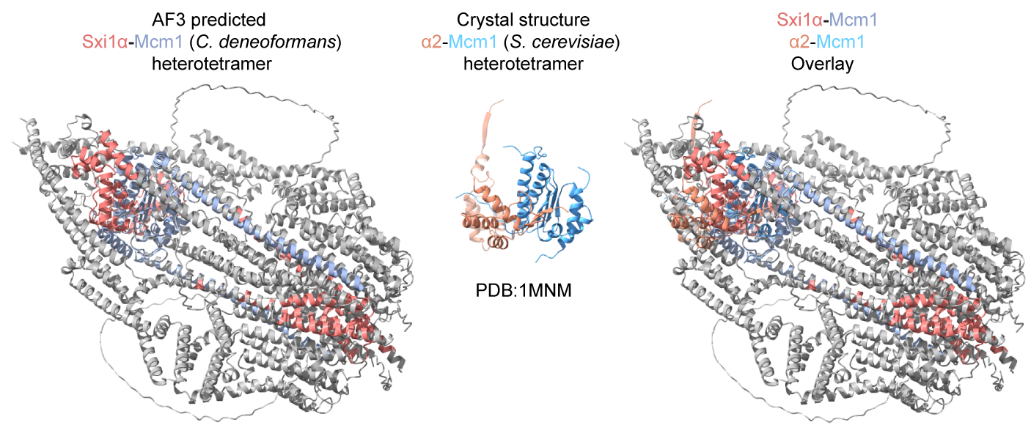

**Supplemental Figure 2. Structural comparison of *C. deneoformans* Sxi1α-Mcm1 and *S. cerevisiae* α2-Mcm1 heterotetramers.**

Left panel: AlphaFold3-predicted structure of the Sxi1α-Mcm1 heterotetramer in *C. deneoformans* (Sxi1α in salmon, Mcm1 in slate blue). Only regions with a predicted local distance difference test (pLDDT) score >60 were colored and surfaced. Middle panel: Crystal structure of the α2-Mcm1 heterotetramer from *S. cerevisiae* (PDB: 1MNM; α2 in orange, Mcm1 in blue). Right panel: Structural overlay of the AF3-predicted Sxi1α-Mcm1 with the α2-Mcm1 crystal structure. Alignment by UCSF Chimera’s MatchMaker yielded a sequence alignment score of 351.9, with an RMSD of 0.432 Å across 76 pruned atom pairs (0.990 Å across all 81 pairs), indicating strong structural similarity between the two heterotetramers.

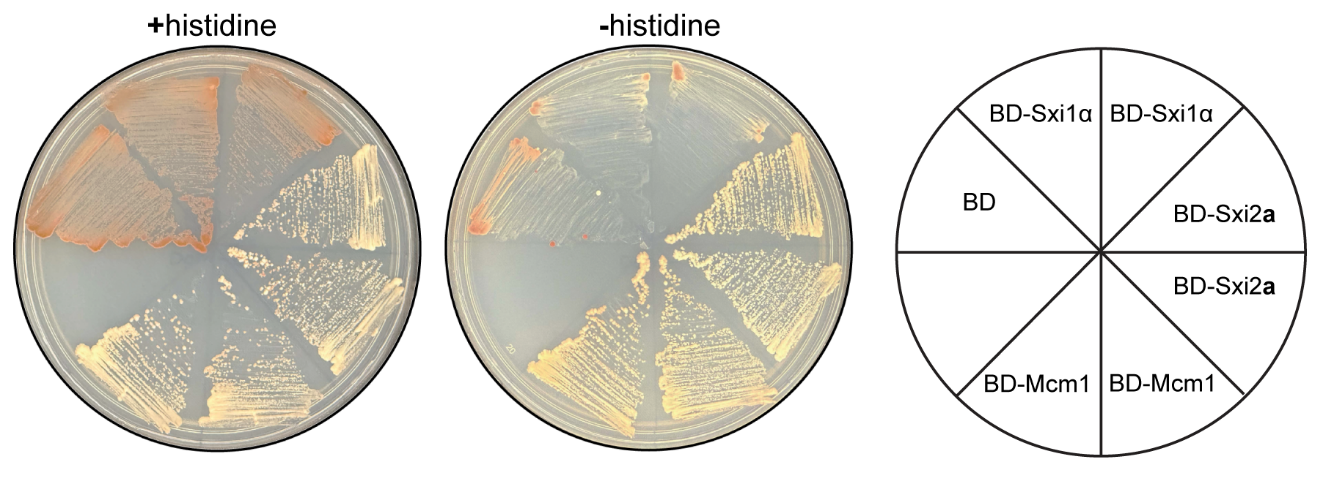

**Supplemental Figure 3. Sxi2a and Mcm1, but not Sxi1α exhibit auto-activation in yeast two-hybrid assays.**

*S. cerevisiae* Y2HGold haploid strains expressing the indicated Gal4 DNA binding domain (BD) fusion proteins were plated on synthetic medium with histidine (+histidine) or lacking histidine (−histidine) to assess activation of the *ADE2 or HIS3* reporter genes*,* respectively. Representative colony color on +histidine medium reflects *ADE2* reporter activity, where reduced red pigment accumulation indicates reporter activation. Two independent clones were tested for BD-Sxi1α**,** BD-Sxi2**a**, and BD-Mcm1. The plate schematic on the right indicates the identity and position of each BD fusion protein.

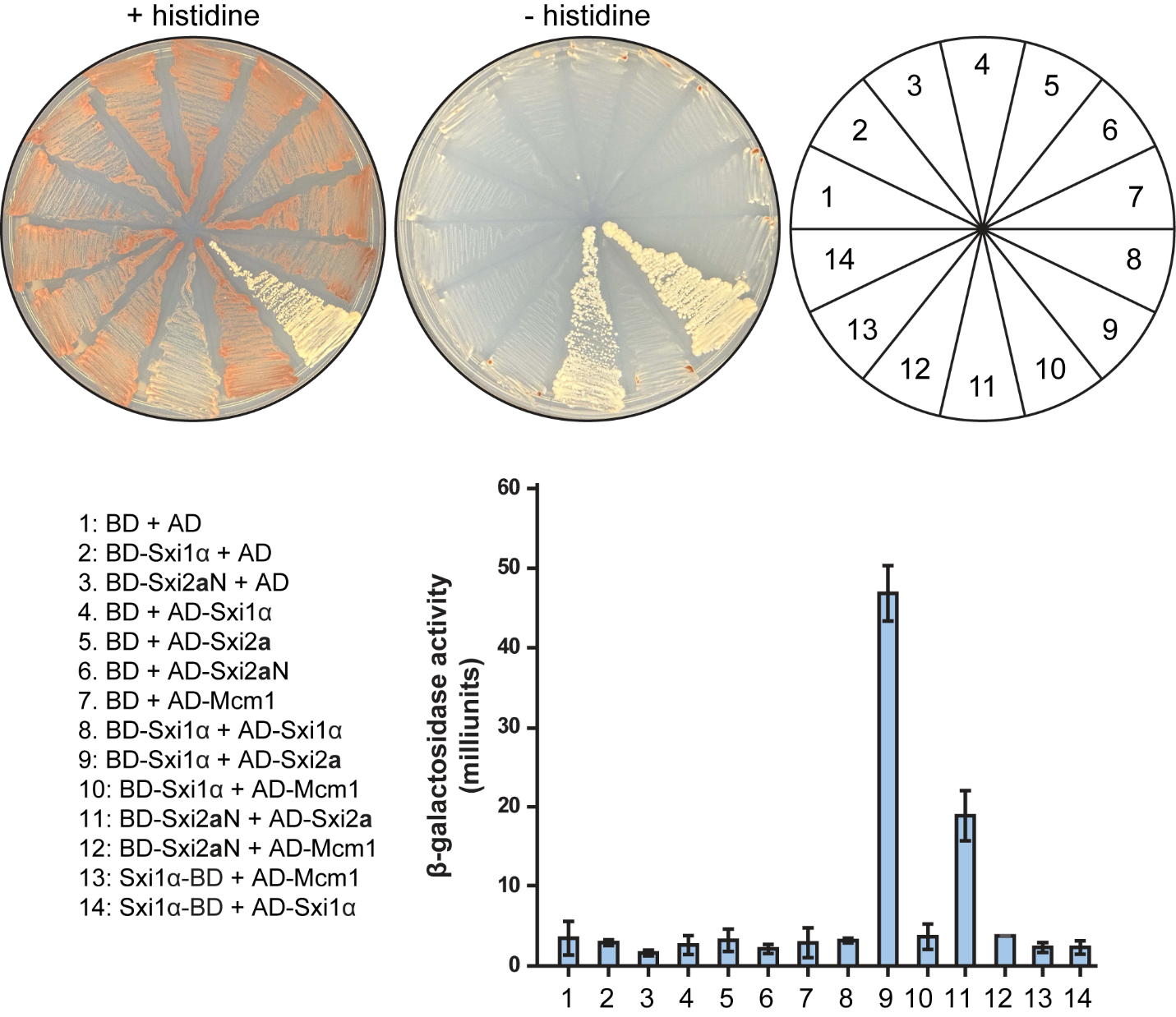
**Supplemental Figure 4. Yeast two hybrid analysis of Sxi1α and Sxi2a interactions.**

*S. cerevisiae* diploid strains co-expressing the indicated Gal4 DNA binding domain (BD) and Gal4 activation domain (AD) fusion proteins were plated on synthetic medium with histidine (+histidine) or lacking histidine (−histidine) to assess activation of *ADE2 or HIS3* reporter genes*,* respectively. Growth on −histidine medium indicates a positive protein–protein interaction. A schematic diagram shows the plate sector numbering (1–14), corresponding to the bait and prey combinations listed. Representative colony color on +histidine medium reflects *ADE2* reporter activity, where reduced red pigment accumulation indicates reporter activation. Quantification of *lacZ* reporter activation was assessed by measuring β-galactosidase activity using a chlorophenol red-β-D-galactopyranoside (CPRG) liquid assay, shown as mean ± standard deviation of three technical replicates.

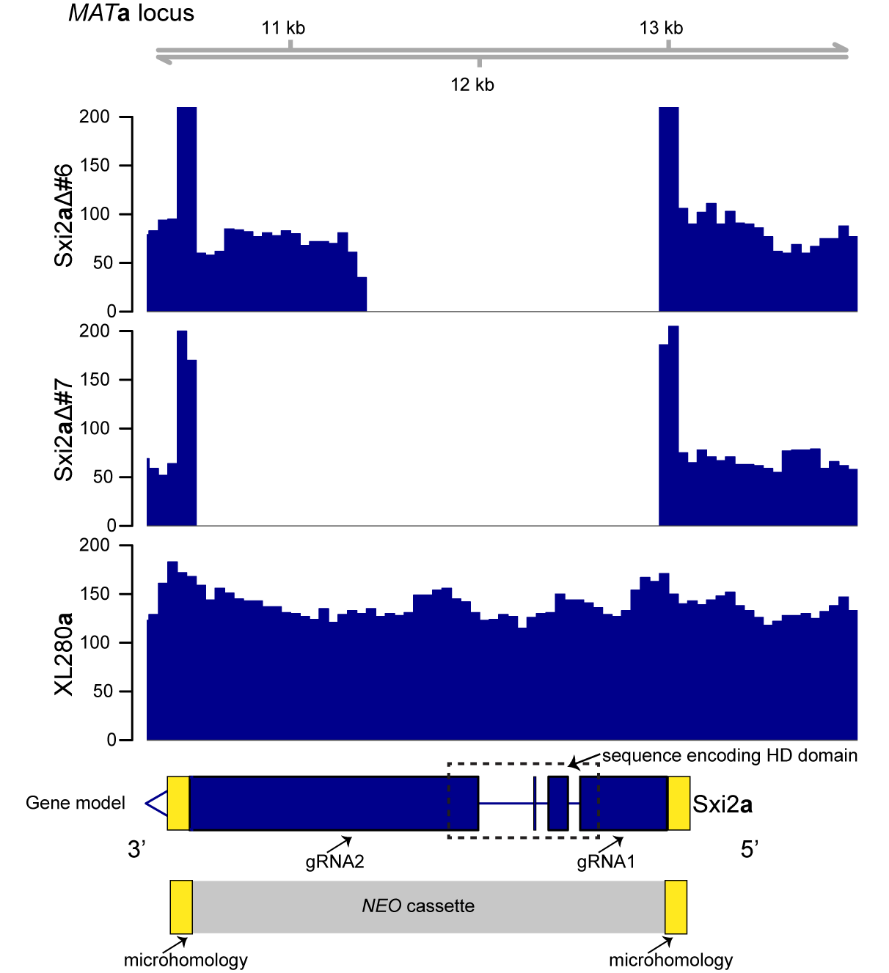

**Supplemental Figure 5. Whole genome sequencing read coverage across the *SXI2a* locus.**

Read depth across the *SXI2****a*** region is shown for the deletion mutants *sxi2***a**Δ#6 and *sxi2***a**Δ#7, with the wildtype XL280**a** strain serving as a control. Coverage gaps indicate regions deleted during mutant construction. The gene model of *SXI2***a** is shown below, with exons represented as blue boxes and introns as connecting lines. Positions of gRNA1 and gRNA2 used for CRISPR mediated deletion are indicated. The DNA sequence encoding the HD domain is indicated by a dashed box. The *NEO* resistance cassette and flanking microhomology arms used for mutant generation are shown in gray and yellow, respectively. The *NEO* resistant cassette is not drawn to scale.

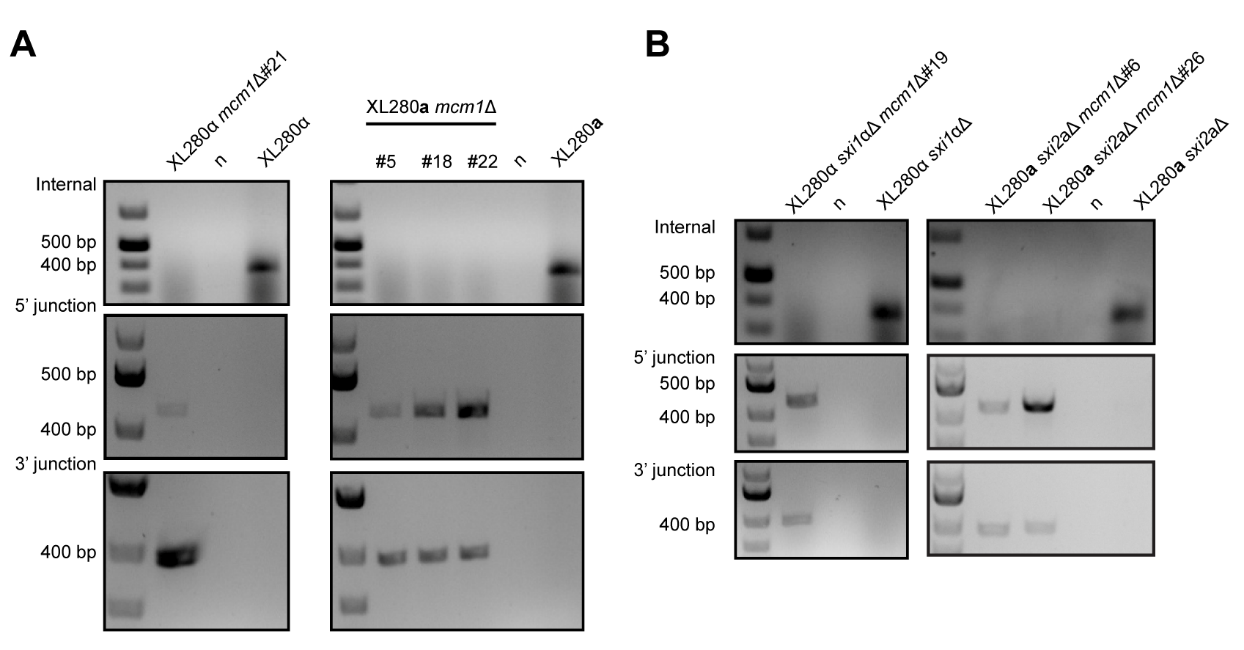

**Supplemental Figure 6. Validation of *MCM1* single and double deletion strains in the XL280 background.**

Generation of independent mcm1Δ (A), sxi1αΔ mcm1Δ and sxi2aΔ mcm1Δ mutants (B). Transformants were verified through internal PCR targeting the ORF, along with 5’ and 3’ junction PCRs specific to the drug resistance marker to confirm proper integration at the genomic locus.

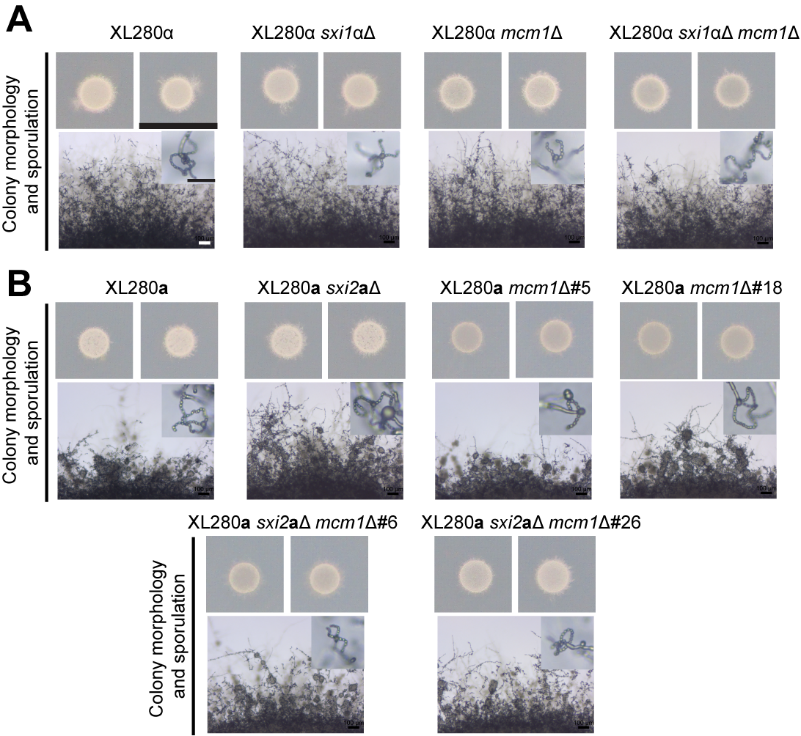

**Supplemental Figure 7. Effects of Mcm1 deletion on unisexual reproduction in the XL280 background.**

Colony morphology (top) and self-filamentation and sporulation phenotypes (bottom) of the independent *mcm1*Δ single deletion mutants (A) and *sxi2***a**Δ *mcm1*Δ and *sxi1*αΔ *mcm1*Δ (B) double deletion mutants grown on MS medium under mating inducing conditions. Scale bars, 1 cm (top panel), 100 μm (self-filamentation) and 20 μm (sporulation) (bottom panel).

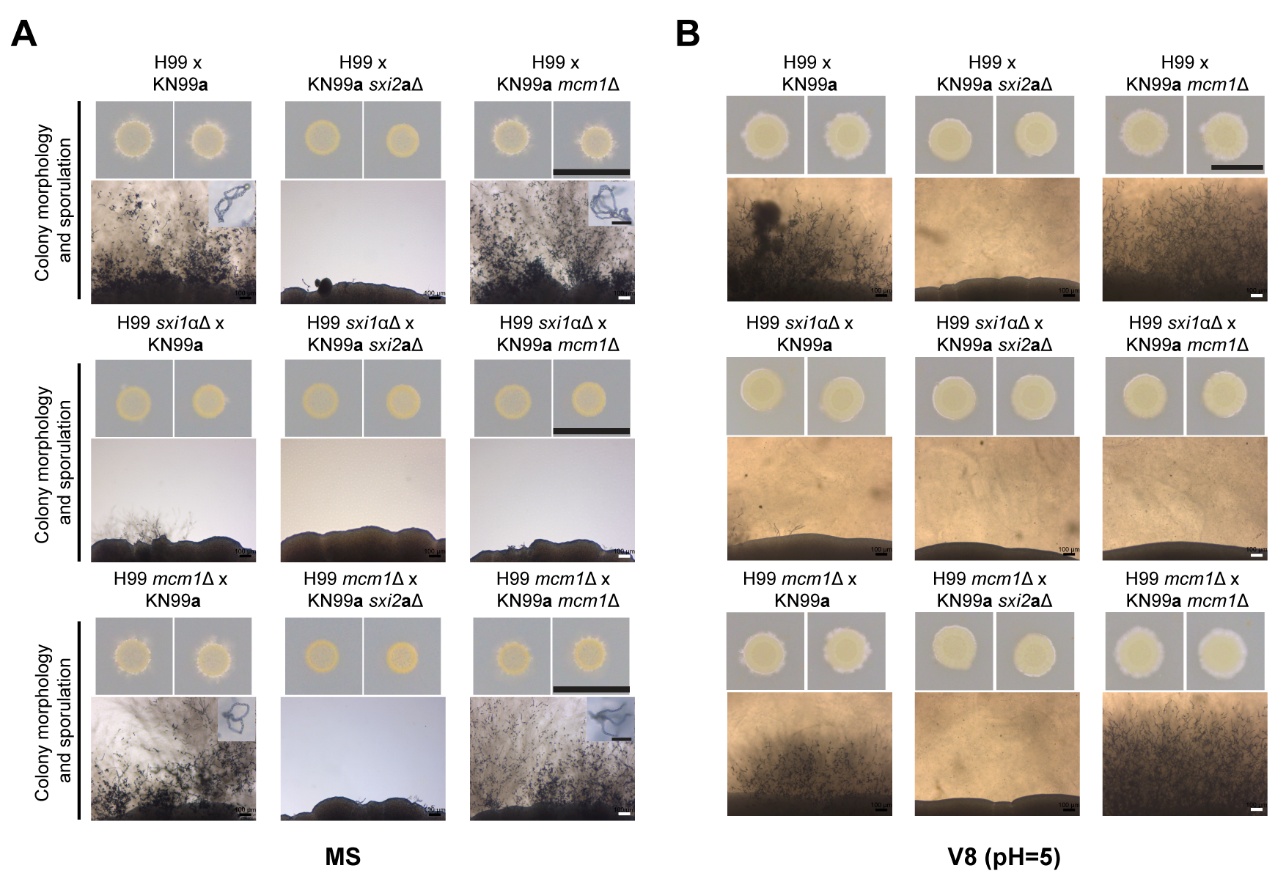

**Supplemental Figure 8. Effects of Mcm1 deletion on α x a sexual reproduction in the H99 background.**

Colony morphology (top) and self-filamentation and sporulation phenotypes (bottom) of the indicated crosses grown on MS medium (A) and V8 (pH=5) (B) under mating inducing conditions. Scale bars, 1 cm (top panel), 100 μm (self-filamentation) and 20 μm (sporulation) (bottom panel).

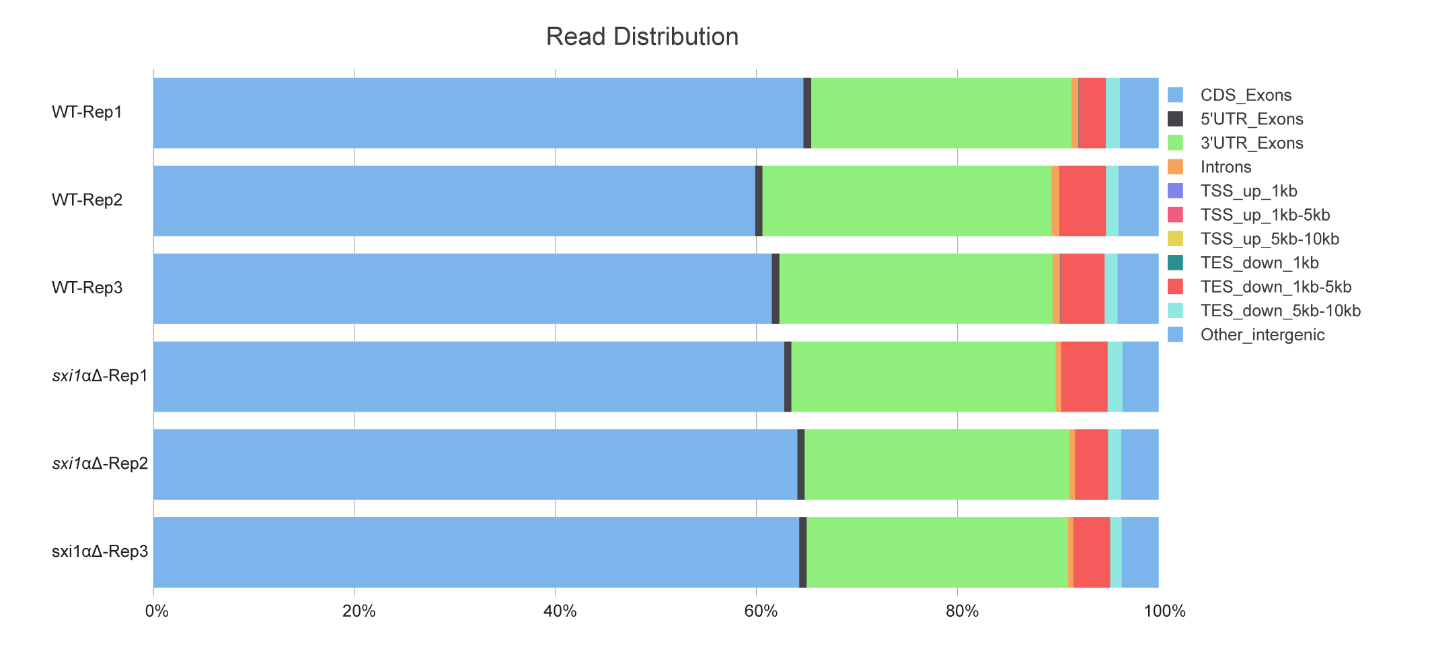

**Supplemental Figure 9.** **Genomic distribution of RNA-sequencing reads across annotated features.** Stacked bar plots show the percentage of RNA-seq mapped reads assigned to different genomic features for wildtype (WT) and *sxi1*αΔ strains across three biological replicates each. Reads were categorized into coding sequence exons CDS exons, 5′ untranslated region exons, 3′ untranslated region exons, introns, regions upstream of transcription start sites TSS at 0–1 kb, 1–5 kb, and 5–10 kb, regions downstream of transcription end sites TES at 0–1 kb, 1–5 kb, and 5–10 kb, and other intergenic regions.

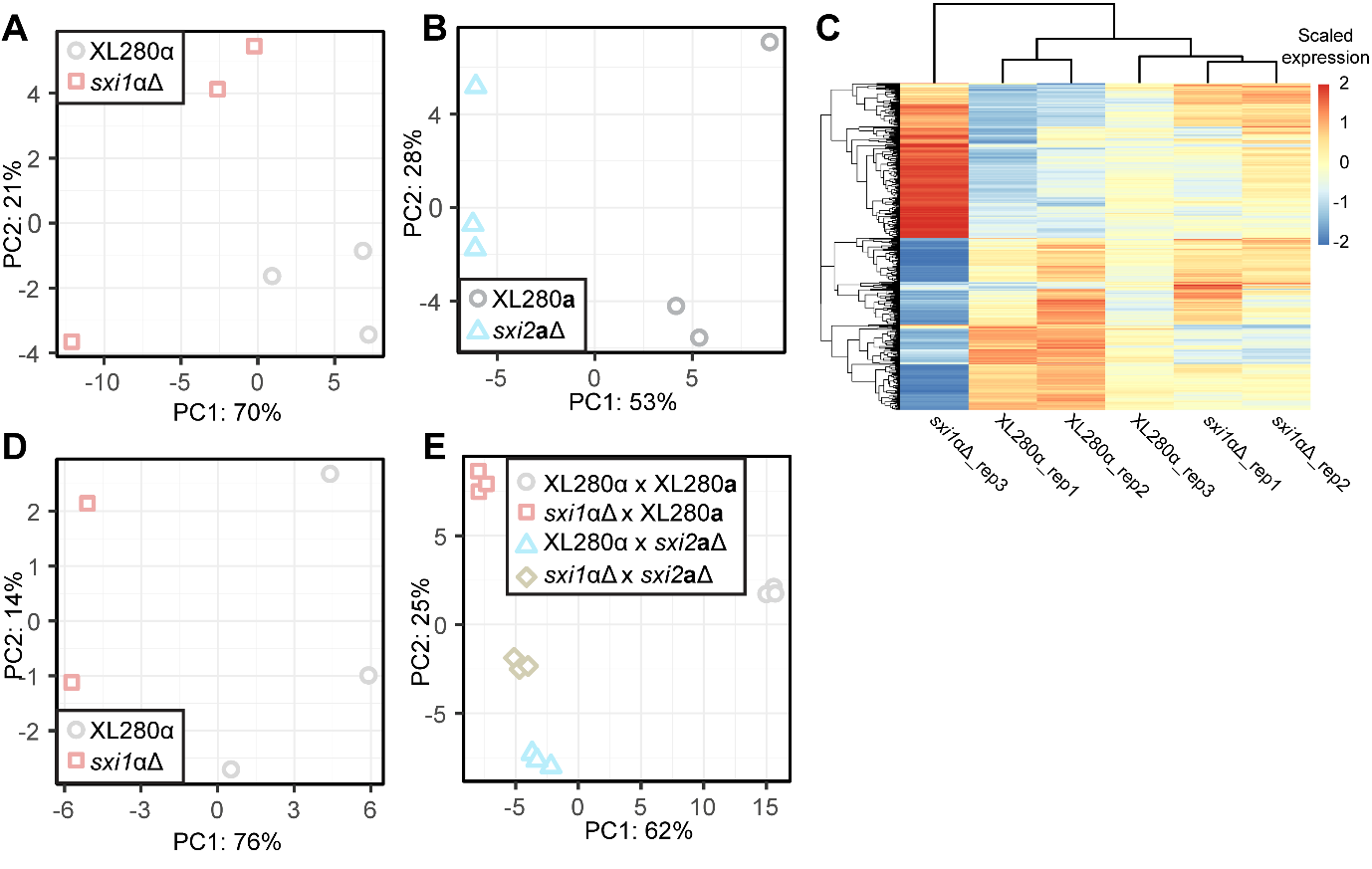

**Supplemental Figure 10. RNA-seq exploratory data analysis.**

All analyses were performed with the top 500 most variable transcripts in each data set. (A and B) PCA of the top 500 most variable transcripts in XL280α (A) and XL280**a** (B) solo-cultures. (C) Hierarchical clustering by sample (x) and transcript (y) of scaled vst-normalized expression values in XL280α and *sxi1*αΔ solo- cultures. (D) PCA of XL280α unisexual crosses excluding *sxi1*αΔ rep 3. (E) PCA of co-culture crosses.

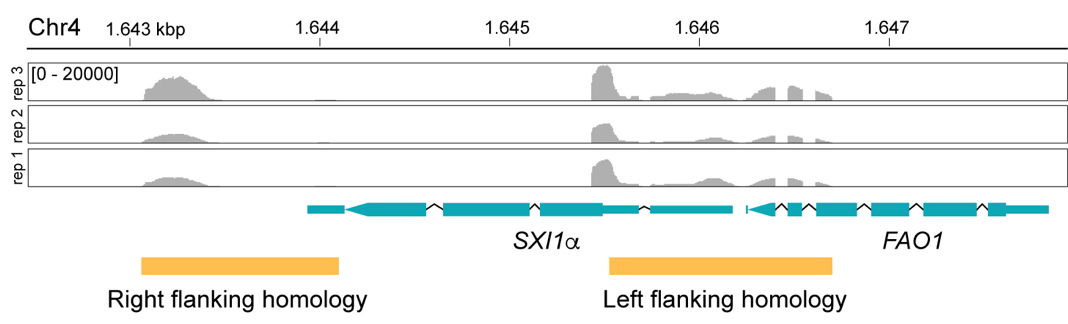

**Supplemental Figure 11.** RNA-seq reads from XL280α *sxi1*Δ unisexual cross mapped to the Sxi1α locus in the XL280α genome. Yellow boxes indicate regions used as homology for the gene deletion cassette.

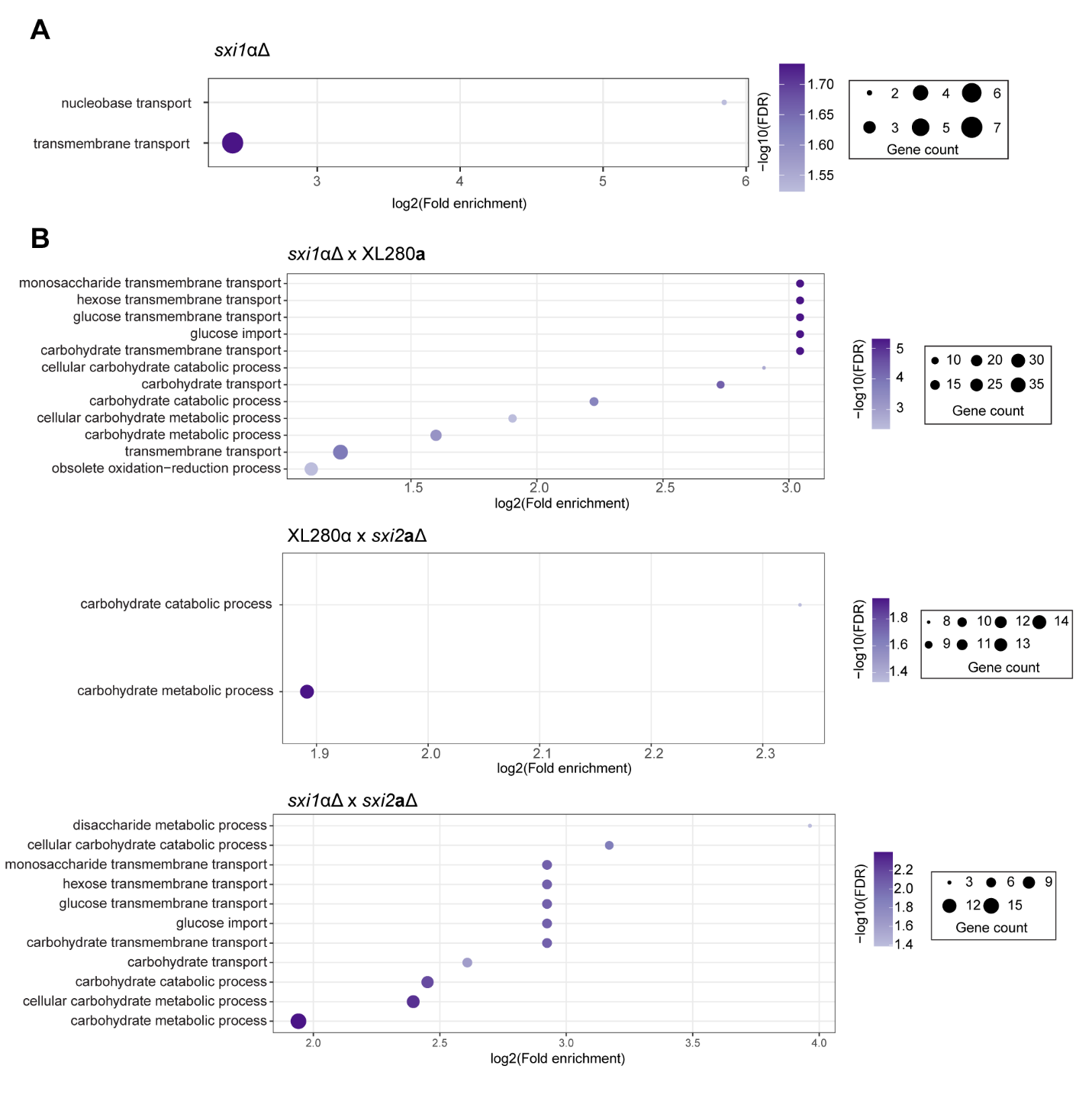
**Supplemental Figure 12. GO term enrichment analysis of DEGs from solo- and co-culture crosses.**

(A) Enriched GO terms in the *sxi1*αΔ solo culture. (B) Enriched GO terms in unilateral (*sxi1*αΔ x XL280**a**, XL280α x *sxi2***a**Δ) and bilateral (*sxi1*αΔ x *sxi2***a**Δ) co-culture mating crosses. Dot size represents the number of DEGs associated with each GO term, and dot color indicates statistical significance as −log₁₀(FDR). GO terms are ranked by log₂(fold enrichment) relative to the whole genome background.

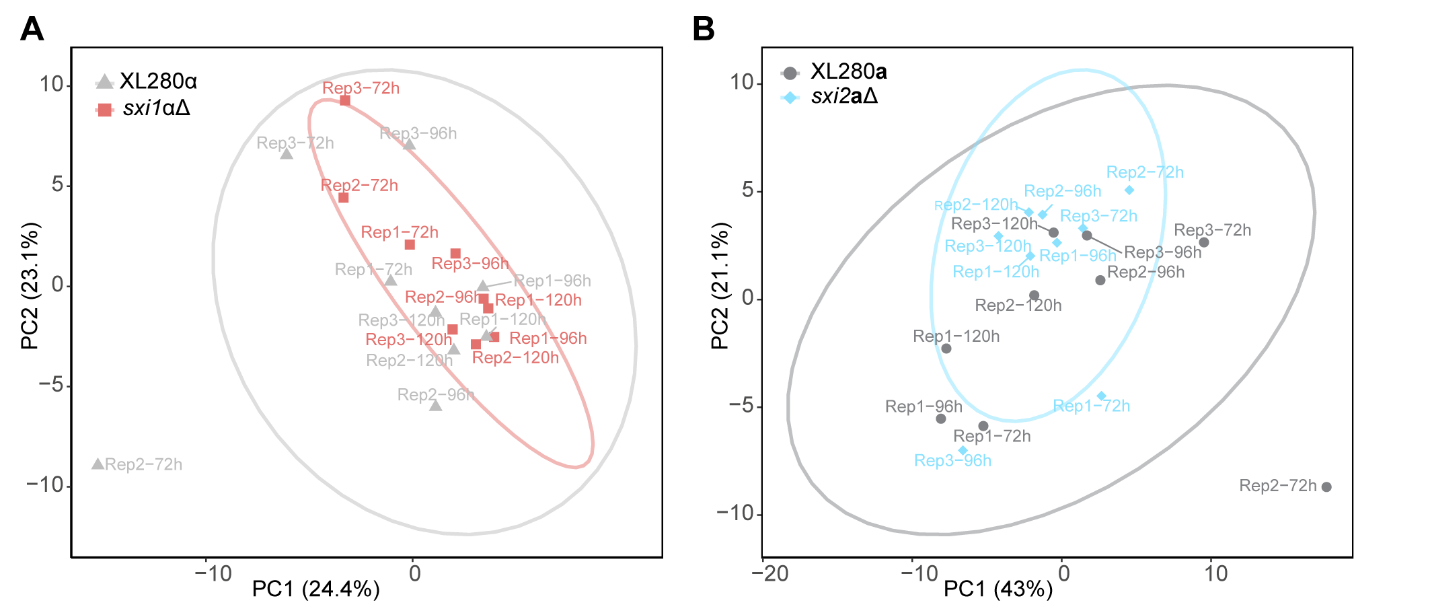

**Supplemental Figure 13. Principal component analysis (PCA) of Biolog YT phenotypic microarray data for wildtype and mutant strains.**

PCA plots show global metabolic activity profiles based on normalized signal values from Biolog YT plates containing 94 carbon sources, measured at 72, 96, and 120 h. (A) *sxi1*αΔ compared to XL280α. PC1 explains 24.4% of the variance, and PC2 explains 23.1%. (B) *sxi2***a**Δ compared to XL280**a**. PC1 explains 43% of the variance, and PC2 explains 21.1%. Each point represents an individual biological replicate, colored by strain, with ellipses indicating the 95% confidence intervals for each group.

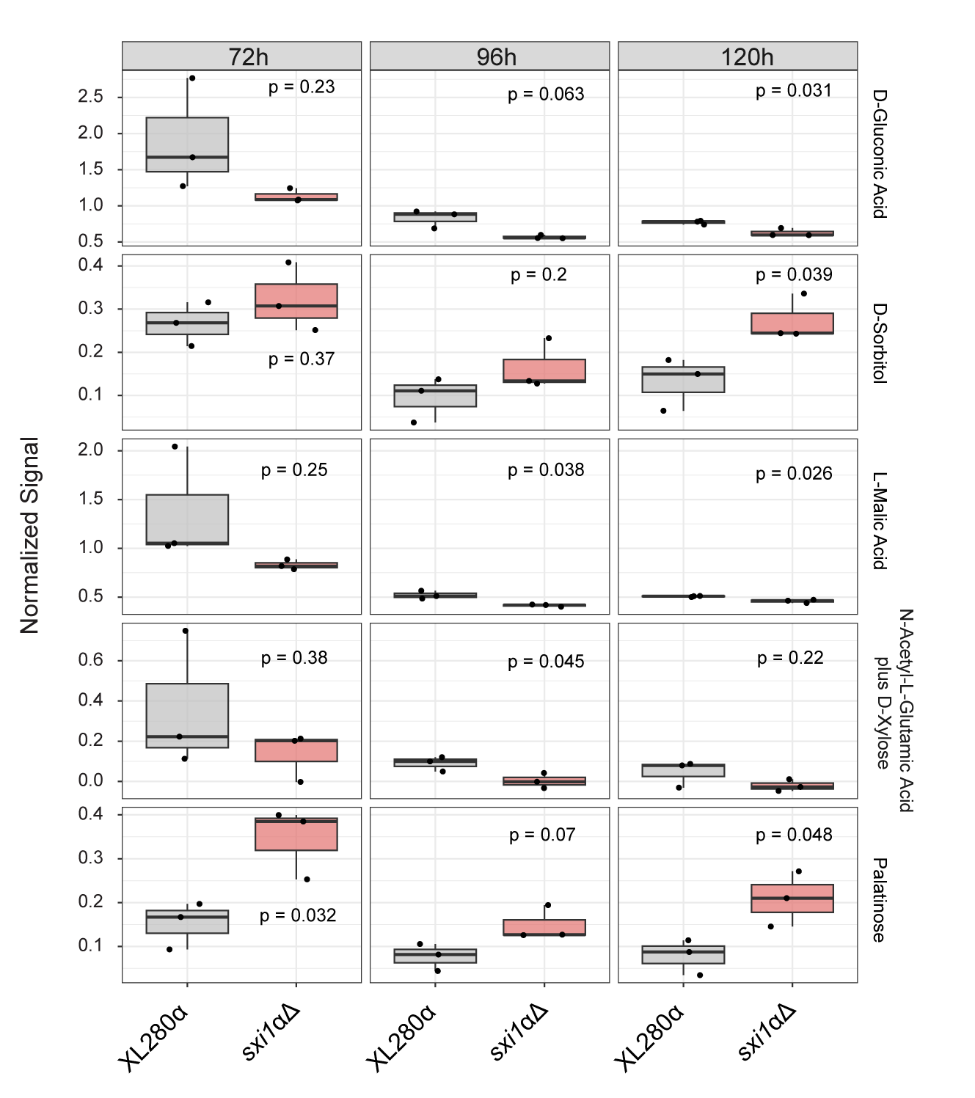

**Supplemental Figure 14. Carbon sources showing significant differences in utilization between sxi1**α**Δ and XL280**α **in Biolog YT phenotypic microarrays.**

Boxplots display normalized signal values for D-gluconic acid, D-sorbitol, L-malic acid, N-acetyl-L-glutamic acid plus D-xylose, and palatinose at 72, 96, and 120 h. Gray boxes represent XL280α and red boxes represent *sxi1*αΔ. p-values were determined by pairwise t-tests comparing mutant and wild type for each substrate at each time point.

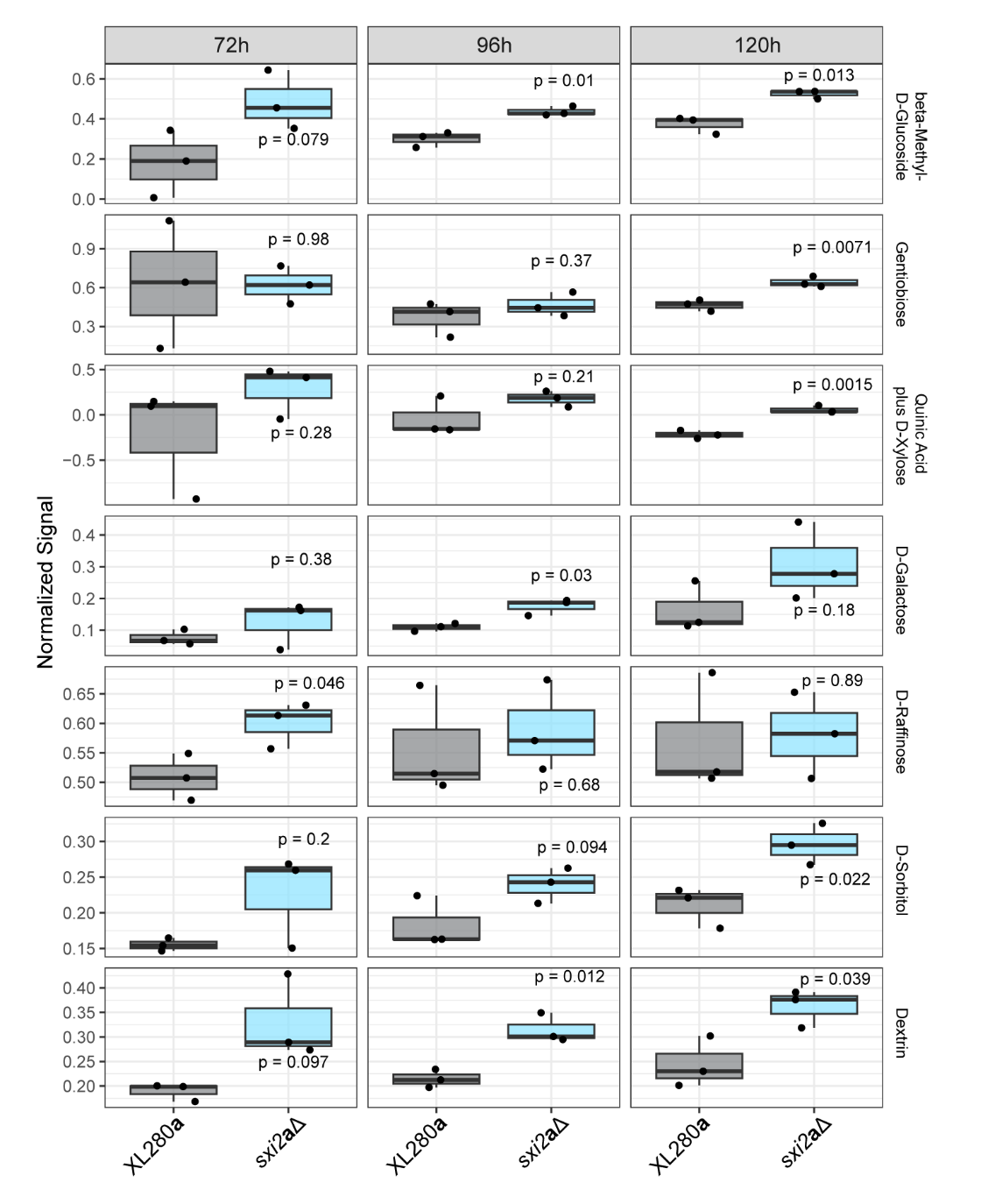

**Supplemental Figure 15. Carbon sources showing significant differences in utilization between *sxi2*aΔ and XL280a in Biolog YT phenotypic microarrays.**

Boxplots display normalized signal values for β-methyl-D-glucoside, gentibiose, quinic acid plus D-xylose, D-galactose, D-raffinose, D-sorbitol, and dextrin at 72, 96, and 120 h. Gray boxes represent XL280**a** and blue boxes represent *sxi2***a**Δ. p-values were determined by pairwise t-tests comparing mutant and wildtype for each substrate at each time point.
